# Abstracted functions for engineering the autonomous growth and formation of patterns

**DOI:** 10.1101/549352

**Authors:** Atri Choksi, Drew Endy

## Abstract

Natural biological patterns arise via the growth, differentiation, death, differential adhesion, communication, and movement of or among cells. Synthetic biologists typically impose explicit genetic control of cell-cell communication and programmable cell state to realize engineered biological patterns. Such engineering approaches do not usually consider the underlying physical properties of individual cells that inevitably contribute to pattern development. To better integrate synthetic genetic systems engineering with natural growth and patterning we derived abstract functions that relate how changes in basic cell properties such as growth rate, length, and radius of curvature result in differences in the curvature, end-point reliability, and texture of borders that define boundaries among growing cell lineages. Each abstracted border function is derived holistically as an emergent consequence of underlying cell physical properties. We experimentally demonstrate control of border curvature to angles of 60° from initial trajectories, control of end-point variability to within 15° of desired target endpoints, and control of border texture between 10 to 60 unit cell lengths. In combination with synthetic genetic control systems, we grow arbitrary two-dimensional patterns including phases of the moon, PacMen, and a yinyang-like pattern. Differences between the idealized and observed behavior of abstracted border functions highlight opportunities for realizing more precise control of growth and form, including better integration of synthetic genetic systems with native cellular properties and processes.

## INTRODUCTION

Patterns are integral to our lives, from chalk drawings made by children to semiconductor chips created via advanced lithography. However, there are three fundamental limitations to how we typically pattern systems today. First, an outside controller must explicitly direct the patterning process. Second, mass production of pattern-based systems often requires expensive, centralized facilities. Third, the so-produced systems are inert and subject to irreversible failures. For example, consider the process and steps leading to a consumer smartphone with a cracked touchscreen. In contrast, living systems routinely realize distributed self-assembly and are capable of self-healing. Better harnessing the ability of biology to grow and pattern should enable new ways of manufacturing complicated systems such as synthetic organs, hybrid biomaterials, and arbitrary patterns (Coore, 1999; Davies and Cachat, 2016; Nagpal, 2001; Teague et al., 2016; Velazquez et al., 2017).

Efforts to engineer synthetic morphogenesis have typically taken two approaches: autonomous pattern formation or autonomous pattern growth. Autonomous pattern formation is the programmed self-organization of existing cells. For example, diffusion and reaction of chemical species was used to realize bullseye patterns, a diffusion based ‘AND’ gate, and an edge detector within lawns of bacteria cells (Basu et al., 2005; Boehm et al., 2018; Kong et al., 2017; Schaerli et al., 2014; Sohka et al., 2009; Tabor et al., 2009). Additionally, differential adhesion was used to implement autonomous phase separation in a clump of cells (Cachat et al., 2016; Glass and Riedel-Kruse, 2018). Alternatively, autonomous pattern growth leverages programmed self-organization of cells as cells grow and divide, with prior work using artificial symmetry breaking, control over cell adhesion, cell-cell signaling, cell proliferation, locomotion, fusion, and death, to facilitate engineering of synthetic morphogenesis (Cachat et al., 2014; Nuñez et al., 2016). Thus far, the most complex patterns realized via growth on a homogenous substrate without external stimuli are sequential or scale-invariant rings (Liu et al., 2011; Payne et al., 2013).

Consideration of how natural biological patterns arise suggests a more complete and holistic approach to patterning that, if applied in combination with synthetic genetic systems, might enable the growth and formation of more sophisticated living patterns. For example, as cells grow and divide, each cell pushes against and buckles in relation to the orientation of its neighboring cells. Mechanical forces produced and experienced by any single cell may be transmitted amongst all cells within a growing ensemble. Modifying any of the physical characteristics of cells such as shape, adhesiveness, or growth rate gives rise to different emergent patterns including furrows, leaf-like shapes, segregated cell lineages, fractal patterns, and heart shaped forms (Korolev et al., 2012; Muller et al., 2014; Odell et al., 1981; Rudge et al., 2013; Steinberg, 1963; Vlad et al., 2014).

While natural patterning systems are wonderful as inspiration, they are also wonderfully complex in that the observed patterns emerge from many compounding mechanisms acting together. For example, growth of Arabidopsis tissue creates mechanical stress that in turn feeds back to amplify local growth rate differences (Uyttewaal et al., 2012). Beyond growth rate and mechanical stress, cell orientation and hormonal signaling contribute to shoot apical meristem patterning (Hamant et al., 2008; Müller and Sheen, 2008). Synthetic approaches might instead allow individual mechanisms to be explored and studied in isolation, allowing for a more complete understanding of the contribution of each mechanism to an overall patterning scheme.

To study such questions and better characterize the limits of current approaches we considered the problem of engineering the growth and formation of arbitrary two-dimensional patterns. As one starting example, we considered if and how a single bacterial cell could be engineered to autonomously grow and divide, eventually producing a multicellular colony manifesting a fully differentiated yinyang pattern. In this case, the specific pattern serves as a heuristic device to test current understanding of morphogenesis and to guide the ability to grow and form synthetic biological patterns. The yinyang pattern itself is not radially symmetric and thus more challenging to realize than previously reported patterns; a single mechanism acting either once or repeatedly over the radius of a colony is insufficient to grow a yinyang. Additionally, the diverse features within a yinyang such as the bend between the white and black halves and the small circles within the halves suggest that multiple patterning mechanisms acting in combination and asymmetrically over multiple length scales may be required. Starting from this initial challenge, we identified the importance of borders in organizing the growth and form of patterns. We sought and were able to develop first-generation mathematical abstractions representing control of border curvature, reliability, and texture as a function of underlying cell properties. We evaluated the extent to which such functions can be practically implemented and developed examples of new forms and patterns that can be realized via a more holistic integration of synthetic genetic programs and natural cellular properties.

## RESULTS

### An initial attempt to grow a yinyang-like pattern

We attempted to engineer a single *E. coli* cell to grow into a yinyang-like colony to explore what opportunities for learning might emerge. A yinyang is composed of a large circle, two embedded circles half the radius of the larger, and two smaller interior circles. The interface between the two mid-sized circles creates a ‘S’-shaped bend. While a two-dimensional ring shape such as a bullseye pattern is radially symmetric and can be collapsed to a one-dimensional square wave along a radial axis in polar coordinates, a yinyang is more complex in that no single one-dimensional pattern can be used to represent its entirety (Figure 1A).

**Figure 1:**
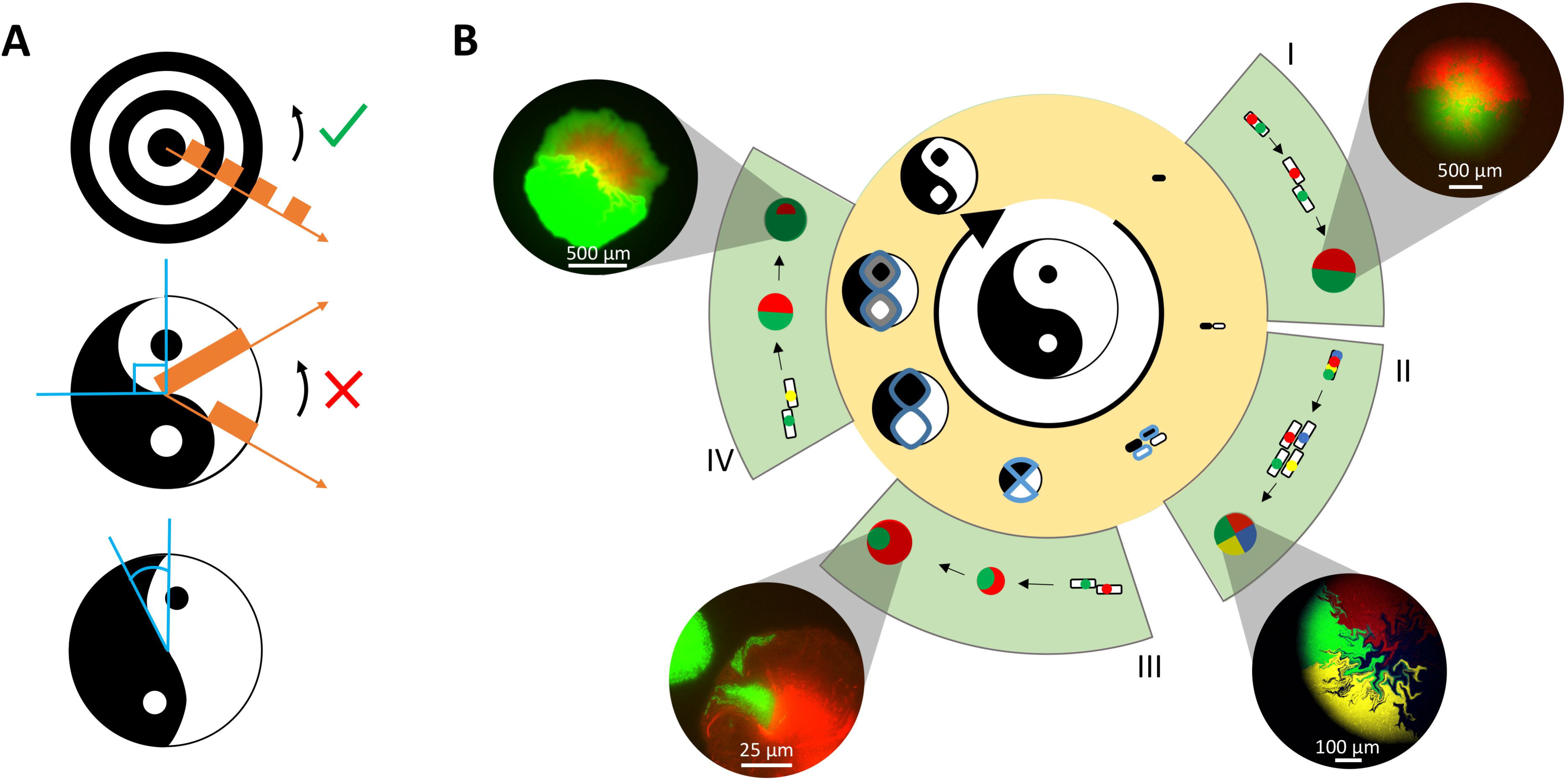
A developmental strategy to grow a yinyang-like colony from a single *E. coli* cell. **(A)** The yinyang pattern is a true two dimensional pattern. In contrast to a bullseye pattern that can be simplified to a square wave in one dimension (top), a yinyang pattern (middle) has no color preserving symmetry. We envision growing a yinyang-like pattern (bottom) in which the curves are more shallow and not as round. **(B)** A proposed mechanism to grow a yinyang-like colony on a homogeneous surface without external stimuli starting from a single *E. coli* cell. The inner white circle shows the desired pattern. The outer gold ring illustrates the hypothesized growth of the yinyang-like pattern with slower growing cells outlined in blue. The outermost green sectors are prototyped, individual patterning steps. (I) Mechanism and implementation of differentiation from a single cell into two and (II) four genetically distinct cell types. Bacterial artificial chromosomes (BAC) with incompatible replication of origins are forced to segregate upon cell division. Images are of colonies grown from a single cell. (III) The four cell types then shape the colony sectors due to differential growth. Image of colony grown from two cells with different growth rates seeded next to each other. (IV) In the last step, encapsulated sectors detect their edge by degrading and sensing AHL. AHL is initially present homogenously in the agar. Cells express an AHL degradation enzyme and express GFP in response to high levels of AHL. As the sector grows larger and AHL is degraded faster, the amount of AHL in the center of the sector decreases. Image of colony grown for 16 hours from two microcolonies where yellow symbolizes both green and red fluorescence.

We strategized a four-step sequential program for directing the growth and formation of a yinyang-like pattern on a homogeneous surface starting from a single cell (Figure 1B). First, a single *E. coli* cell differentiates into two cells of different states, forming the white and black halves of the yinyang (Figure 1B.I). Second, each color-unique lineage differentiates again into fast and slow growing subpopulations (Figure 1B.II). Third, differences in growth rate result in a ‘S’-shaped bend between the white and black halves (Figure 1B.III). Finally, a synthetic genetic edge detection program specific to each of the slower growing cell subpopulations produces a second cell state differentiation event and color reversion within the “eyespots” of each yinyang half (Figure 1B.IV).

We empirically prototyped each of the four steps as envisioned. First, we used plasmid segregation to achieve one and two differentiation events (Nuñez et al., 2016). Approximately one-third of colonies grown from single cells containing two plasmids resulted in bipartite colonies, signifying a single differentiation event occurring early on during colony growth (Figure 1B.I). Second, we used four-plasmid segregation to engineer two differentiation events that result in colonies with four quarters, however only rarely (∼0.5% of the time) (Figure 1B.II). Third, we realized curved borders by using growth rate differences between neighboring seed cells (Figure 1B.III); we note that curves realized via growth rate differences alone are expected to be less stiff than what is found in a yinyang (Figure S1). Fourth, we prototyped a sector edge detector composed of nine synthetic genetic devices (Figures 1B.IV and S2). Cell sectors harboring the edge detector fluoresced green at their edges with an edge width of approximately 150 µm. Each of the four individual steps behaved as expected, to a degree, but with significant differences from the ideal desired behavior (below).

### Borders are rough and curve unreliably under standard conditions

The individual steps used in prototyping the growth and formation of a yinyang revealed more complexity in patterning then we desired. For example, often the same initial conditions resulted in different specific patterns. For example, plasmid segregation resulted in some colonies that were bipartite, but also resulted in colonies with no sectors or many sectors (Figure S3). As a second example, although the average trajectory of any given border might realize a desired curvature, closer examination of borders revealed cells protruding into adjacent cell lineages, creating a unique roughness or texture to any specific border (Figure S4).

We therefore sought to more formally explore factors contributing to border curvature, roughness, and reliability using modeling and simulation. In a typical simulation, we started by modeling two rod shaped cells with radii of 0.5 µm and length of 2.5 µm seeded symmetrically next to each other; growth, division, and buckling of seed cells and their offspring were simulated against a fixed wall (Figure S5) (Rudge et al., 2012). We observed how macroscopic border characteristics arise as a function of individual cell properties, and also how these characteristics vary from one simulation run to the next; individual simulations vary from one another due to random differences in cell length at division and exact buckling angle (Supplementary Materials). For example, we observed that a border formed by two adjacent populations growing at equal rates against a fixed wall is expected, on average, to form a line perpendicular to the fixed wall but that individual simulated colonies produce borders that vary in end point angles according to an apparent Gaussian distribution (Shapiro-Wilk test, p = 0.0073) with a standard deviation of 16.6° (Figures 2A and S7).

**Figure 2:**
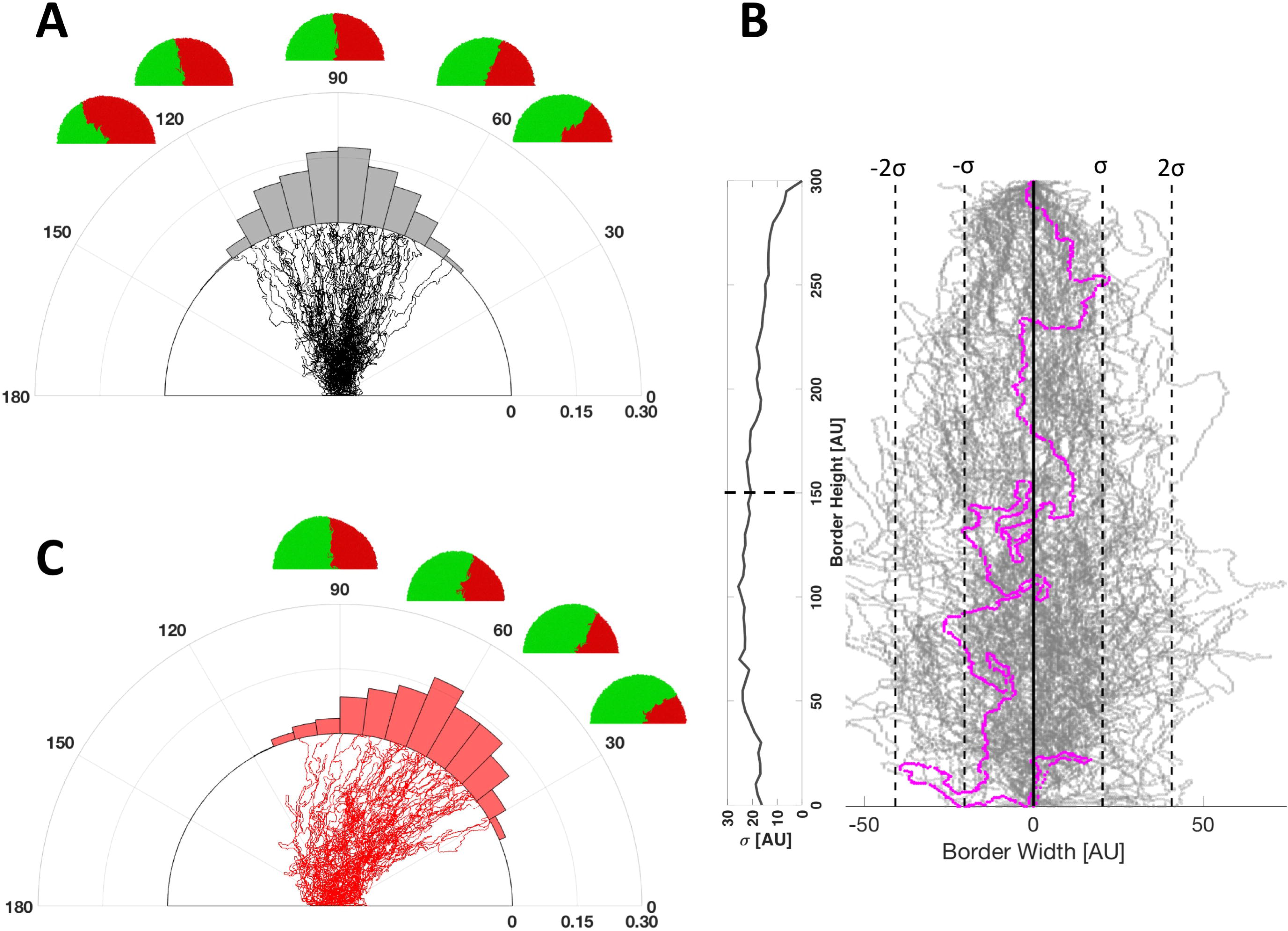
Borders between cell lineages are rough and unreliably curve due to variation in cell length at time of division and random cell buckling events. **(A)** Distribution of border endpoints from repeated simulations of red and green cells growing adjacent to each other at identical growth rates. Sample simulation results are above the polar histogram and sample borders are plotted below. N = 1000. **(B)** Border endpoints are aligned across 100 simulations and a single border is highlighted in pink. Standard deviation of border width is calculated along border height. Standard deviation of border width at 150 AU is plotted as a dotted line. **(C)** Borders unreliably curve under standard conditions. Border endpoint distribution when the growth rate ratio of red cells to green cells is 0.875. N = 1000.

We next explored expected differences in border roughness and texture by aligning simulated border endpoints and quantifying microscopic differences in individual borders compared to an idealized average, straight-line border. We found borders were expected to protrude up to a third of their length away from the expected normal axis (Figure 2B). Finally, we explored how differences in growth rate can result in curved borders. For example, when the growth rate ratio of adjacent seed cells was 0.875 we observed simulated border curves with a distribution mean endpoint angle of 64° and standard deviation of 18°±0.8° (Figure 2C).

Given that patterns are defined in part via the borders that arise via underlying physical properties among populations of interacting and growing cell lineages, but that the exact relationships between individual properties and resulting borders is complicated, we sought to better understand how basic border characteristics arise as a function of changing cell properties. More specifically, we sought to understand and abstract the relationships between individual cell properties and control of border curvature, texture, and end-point reliability.

### Abstracted functions to engineer curvature, texture, and reliability of borders

Border features can arise holistically from a diversity of cell physical properties (Figure 3A-C) (Korolev et al., 2012; Muller et al., 2014; Odell et al., 1981; Rudge et al., 2013; Steinberg, 1963; Vlad et al., 2014). We more formally explored how to control border curvature, texture, and reliability via high-throughput simulation (Figure 3A-C). We adapted the modeling and simulation approach used in our pilot explorations for high-throughput cloud computation platforms such that many of thousands of simulations representing independent tuning of cell physical properties could be realized (Supplementary Methods). From these simulation results we abstracted approximate forms for functions representing the curvature, texture, and endpoint reliability of borders as a function of cell radius of curvature, cell length, cell growth rate, and drag coefficient per cell length, resulting in our development of the following abstracted functions (Figure 3D-F):

**Figure 3:**
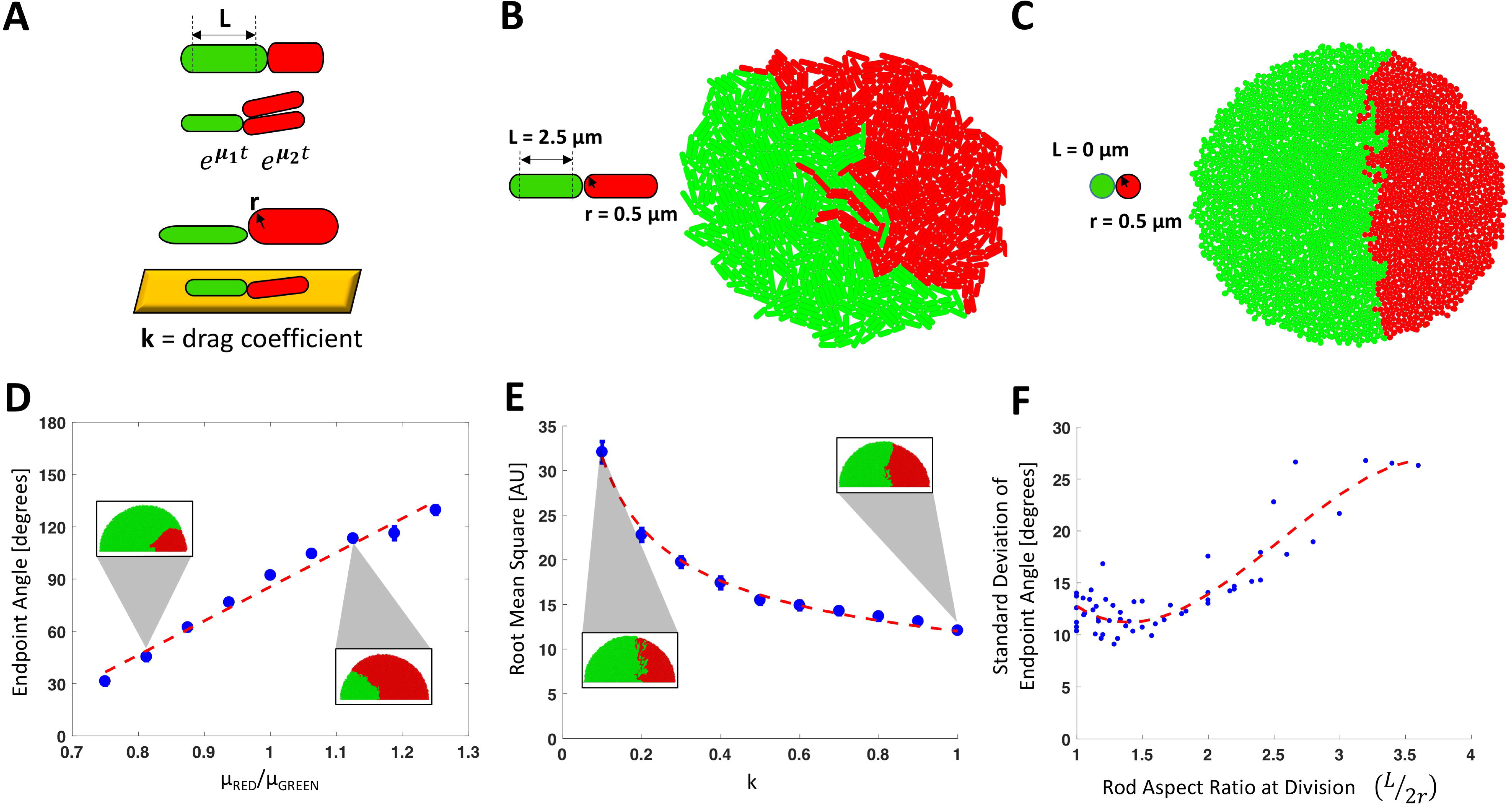
Cell length, radius of curvature, growth rate, and substrate drag coefficient can be abstracted into border functions defining border endpoint angle, roughness, and reliability. **(A)** Border features arise holistically from a diversity of cell physical properties. Physical characteristics at the cellular level such as cell radius of curvature r, cell length L, drag coefficient per cell length k, and cell growth rate µ, can give rise to border features. Colonies grown from cells with cell length of 2.5 µm and a radius of curvature of 0.5 µm and cells with cell length of 1.0 µm and a radius of curvature of 0.5 µm. **(D)** The curvature border function. Plot of mean and standard deviation of border endpoints with different growth rate ratios. Sample simulations in which growth rate ratios of red to green cells are 0.8125 and 1.125 are shown. Red dashed line is linear fit of endpoint angles ranging from 30 to 120 degrees R^2^= 0.9772. N = 100. **(E)** The texture border function. Plot of the average root mean square of borders from average border trajectory at different drag coefficients per cell length. Sample simulations in which drag coefficients are 0.1 and 1 are shown. Red dashed line is a best fit power function R^2^= 0.9941. N = 100. **(F)** The reliability border function. Plot of the standard deviation of border endpoint angles at different rod aspect ratios at division (cell length over cell diameter at time of division). A rod aspect ratio of 1 translates to cells that divide into spherical daughter cells. A third degree polynomial fit R^2^= 0.8268. N = 100.

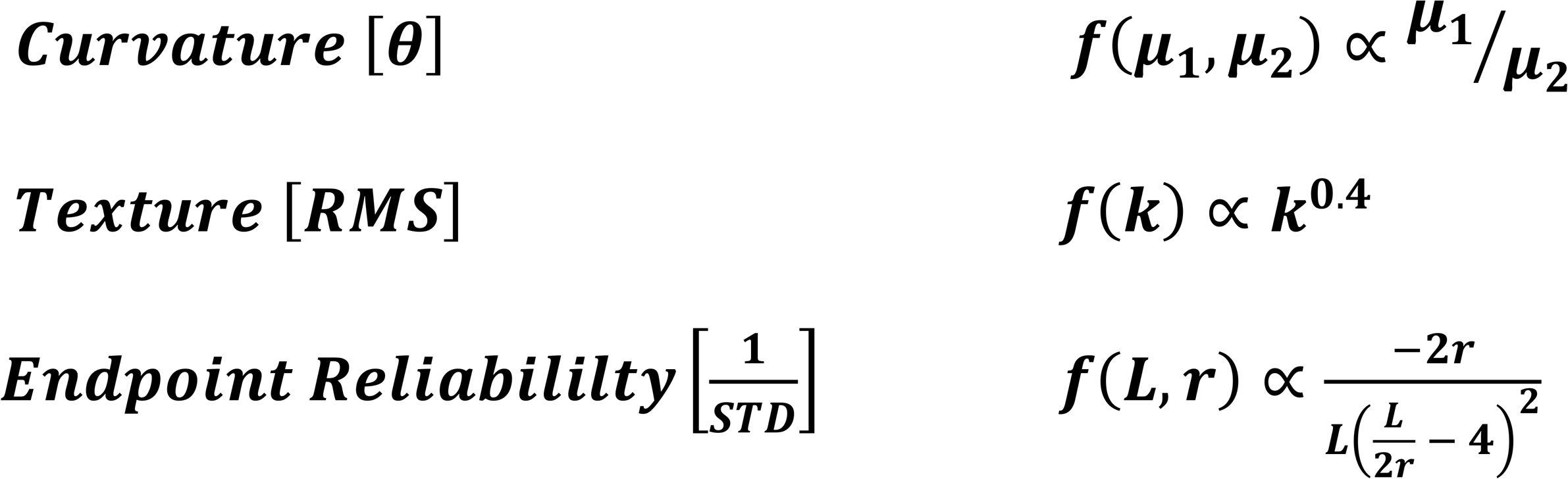

where ***r*** is cell radius of curvature, ***L*** is cell length, ***µ*** is cell growth rate, and ***k*** is drag coefficient per cell length. We observed that the curvature border function is expected to depend linearly with the ratio of cell growth rates between a border endpoint angle range of 30° to 120° and is independent of cell shape and drag coefficient (Figure 3D, S6C and S8A). We next observed that border texture can be controlled independent of border endpoint reliability by modulating the drag coefficient of cells (Figure S8B-C). Additionally, we observed the expected root mean square difference of a border from its average trajectory increases as a power law from ∼12 AU to ∼32 AU as the drag coefficient of cells decreases (Figure 3E). We reconfirmed that borders should be expected to achieve a specific endpoint angle less reliably as cells become more rod-like (Rudge et al., 2013), with endpoint angle standard deviation increasing linearly from 10° to 27° (Figure 3F and S8D-F). We noted, unexpectedly, that spherical cells should not grow into the smoothest or most reliable borders. Instead, we found that very short rod-shaped cells (1.3 aspect ratio at division) are expected to form the most reliable and smoothest borders (standard deviation of border endpoint angles of ∼9°).

### Spherical cells should grow patterns reliably, smoothly, and modulate border curvature

A more detailed comparison of simulated borders formed from rod-shaped versus spherical cells revealed the extent to which spherical cells are expected to grow patterns more reliably than rod shaped cells (Figure 4A), with end point angle distribution standard deviations of 11.4° and 16.6° for spherical and rod-shaped cells, respectively. Spherical cells also formed simulated patterns more smoothly than rod shaped cells, with aligned borders protruding only one-sixth of their height away from the normal axis (Figure 4B). Finally, a comparison of border curvatures realized by tuning cell growth rates suggests that spherical cells achieve similar endpoints on average as rod shaped cells (Figure 4C-E) but that spherical cells achieve the same mean border endpoints with, on average, half the standard deviation as rod shaped cells.

**Figure 4:**
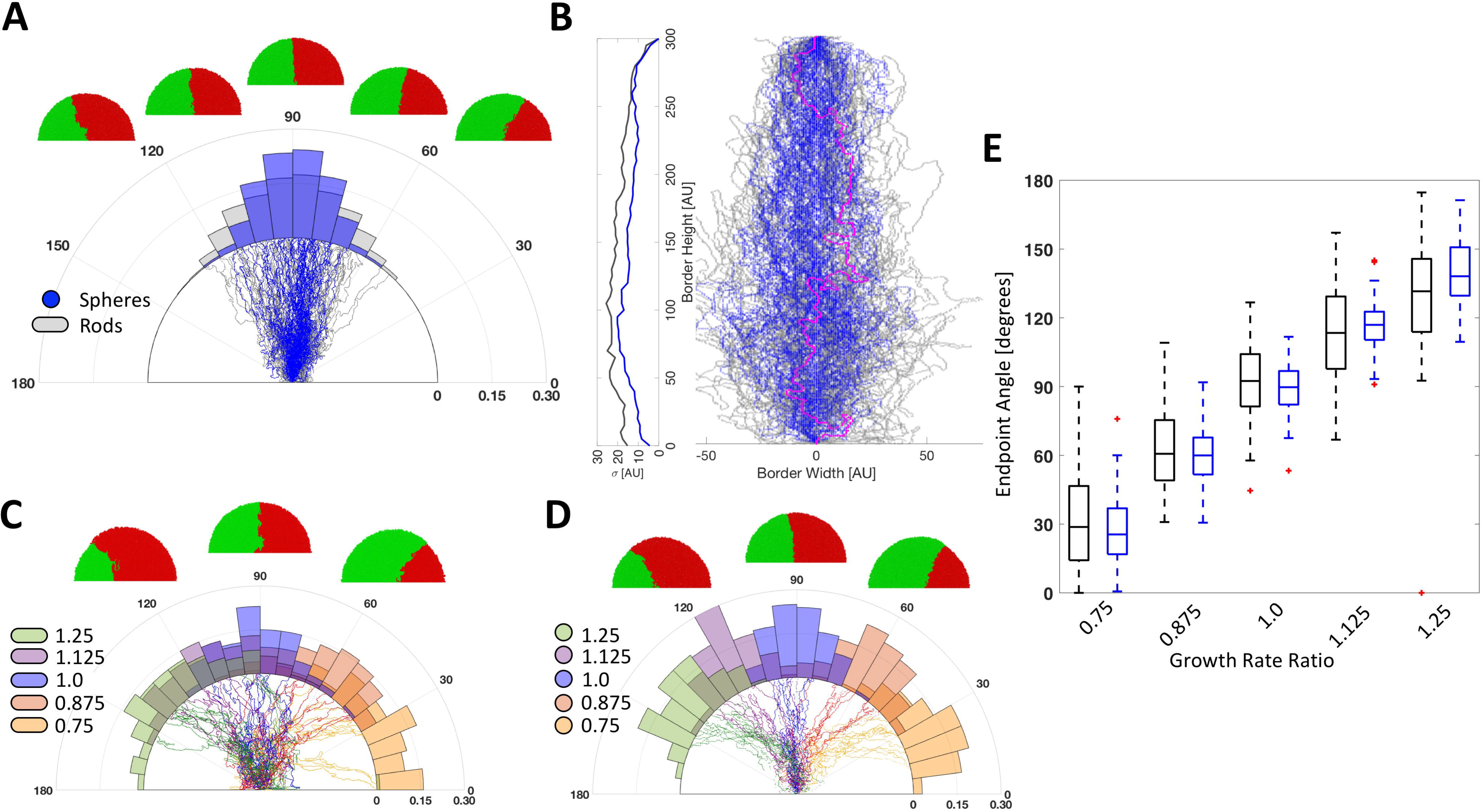
Spherical cells are predicted to make smoother and more reliable borders compared to rod shaped cells. **(A)** Distribution of border endpoints from repeated simulations with spherical cells (blue) compared to rod shaped cells (gray). N = 1000. **(B)** Comparison of aligned borders from spherical cells (blue) and rod shaped cells (gray). A single border simulated with spherical cells is highlighted in red. **(C)** Border endpoint distributions when the red over green cell growth rate ratios are 1.25, 1.125, 1.0, 0.875, and 0.75 with rod shaped cells and **(D)** spherically shaped cells. N = 100. **(E)** Box plot of border endpoint distributions of rod shape (black) and spherical (blue) cells grown at different growth rate ratios.

### Spherical cells grow patterns reliably, smoothly, and modulate border curvature

We tested both our abstracted border functions and our practical ability to implement control over border curvature, texture, and reliability by experiment. We adapted and established a method to micromanipulate two different cells types next to each other for consistent cell seeding (Figure S10-11). We used KJB24 cells, which are spherical *E. coli* cells that are ∼0.7 µm in radius, and DH5*α*Z1 cells, which are rod shaped *E. coli* cells 3 µm in length and ∼0.7 µm in radius (Begg and Donachie, 1998; de Pedro et al., 2001). As predicted, spherical cells grew patterns more reliably (Figure 5A); while rod shaped cells had a distribution of border endpoints with a standard deviation of 22.7°,+3.6°, –2.4°, spherical cell border endpoints varied by a smaller standard deviation of 15°,+2.4°, –1.6°. Aligned borders from colonies consisting of rod shaped cells varied from the normal axis by as much as 60 µm while spherical cells only varied by about 10 µm or less (Fig 5B, C, and F). Border curvatures were achieved by tuning growth rates with bacteriostatic antibiotics without significantly modifying cell shape (Figure S12-14) (Ciak and Hahn, 1958; Garrett and Miller, 1965). As before, spherical cells more reliably achieved border curvatures with an improvement in standard deviation of 7° (Figure 5D-E).

**Figure 5:**
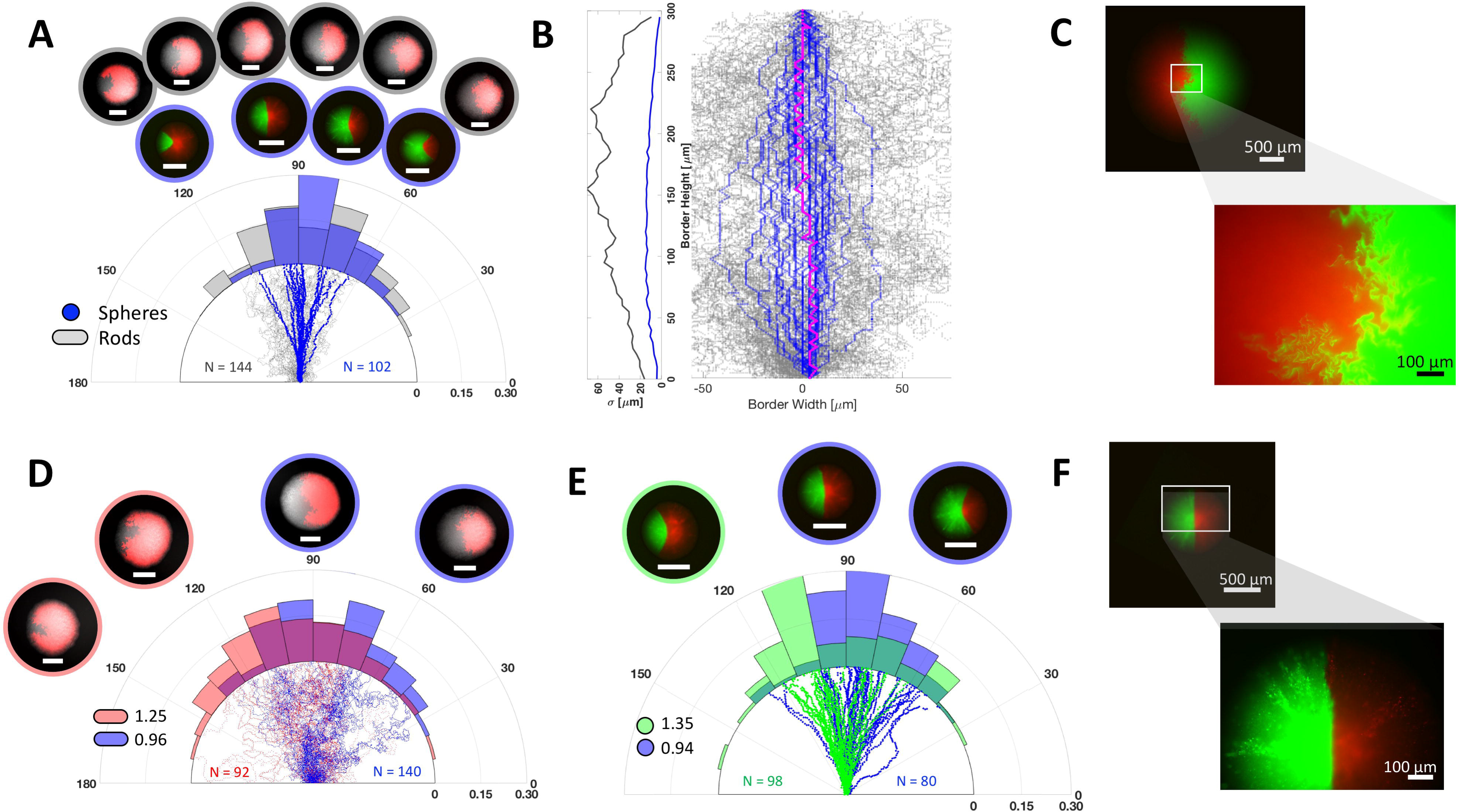
Experiments show that spherical cells make smoother and more reliable borders compared to rod shaped cells. **(A)** Distribution of border endpoints from grown microcolonies of rod shaped cells (gray) and spherical cells (blue) with sample colony images above the histogram and sample traces below. **(B)** Comparison of aligned borders of rod shaped cells (gray) and spherical cells (blue) with standard deviations calculated over every 5 µm. A single border grown from spherical cells is highlighted in pink. **(C)** 2X and 10X Images of a colony grown from rod shaped RFP and GFP expressing microcolonies seeded next to each other. **(D)** Distribution of border endpoints with the growth rate of green cells slowed with 0 µg/mL and 0.6 µg/mL Chloramphenicol. Cells fluorescing red carry Chloramphenicol resistance. **(E)** Distribution of border endpoints with the growth rate of green cells slowed with 0.1 µg/mL and 0.4 µg/mL Tetracycline. Scale bars 500 µm. **(F)** 2X and 10X images of colony grown from spherical cells.

### Combining holistic patterning and explicit genetic systems for the growth and formation of arbitrary two-dimensional patterns

Abstracted border functions for programming pattern growth should ideally be useful for engineering more complex arbitrary two-dimensional patterns in combination with synthetic genetic systems. We thus engineered the growth and formation of a variety of biological patterns arising from seed patterns comprised of two or more cell types that combine differences in cell growth rates as well as synthetic genetic programs (Figure S15-16). For example, we grew patterns that appear as different phases of the moon starting from two cell types (Figure 6A). One cell type harbors a sector edge detector plasmid and is resistant to tetracycline while the other cell type only expresses green fluorescent protein (GFP). Programmed decreases in the growth rate of GFP-expressing cells via addition of tetracycline resulted in waxing phases of the moon (Figure 6A). As a second example, we engineered “PacMen” with varying mouth sizes by seeding an additional cell type that weakly expresses red fluorescent protein (RFP) (Figure 6B). Lastly, we engineered a yinyang-like pattern by seeding an additional sector edge detector cell-type that fluoresced red at its edges (Figure 6C, S17, and S18).

**Figure 6:**
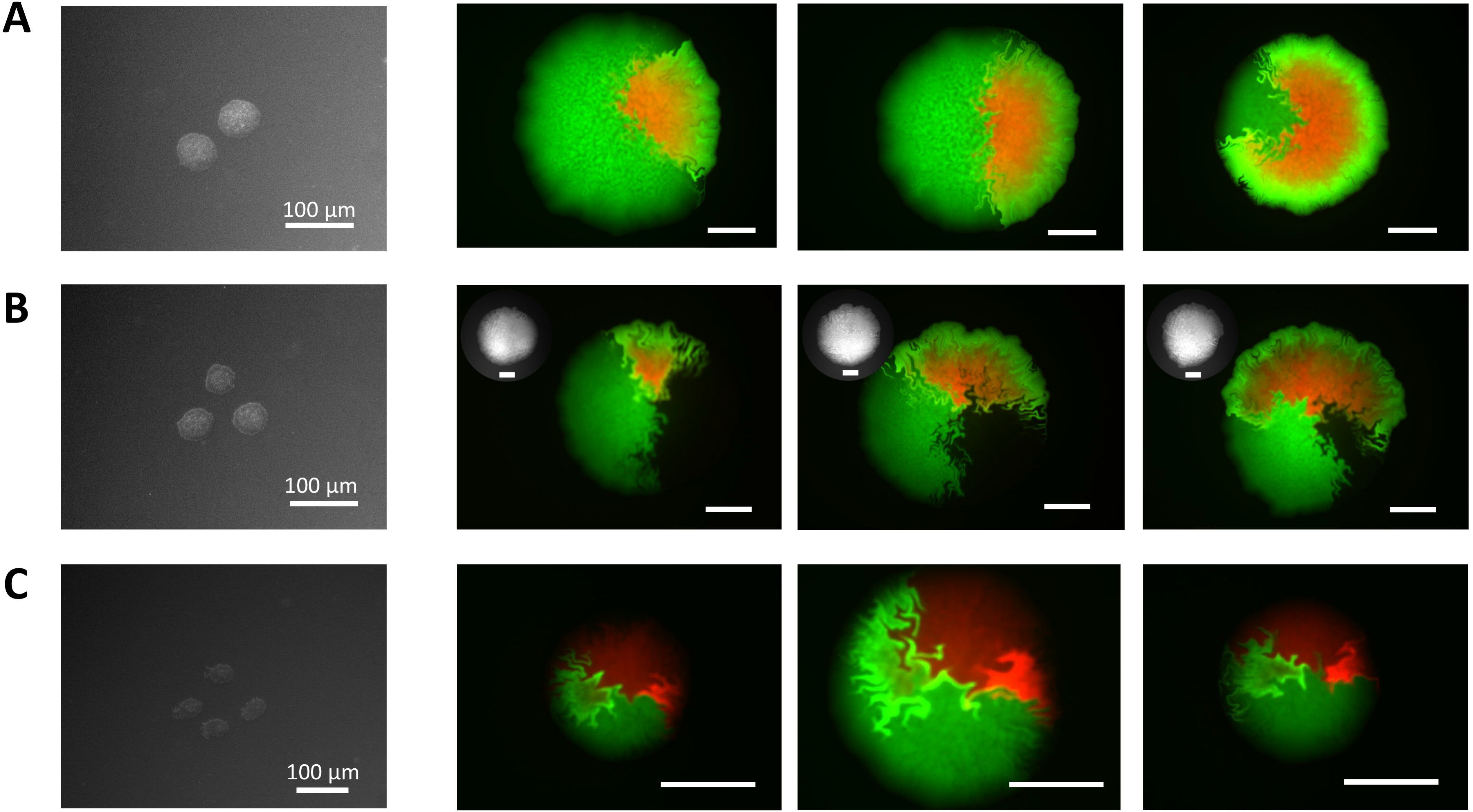
Growth of arbitrary synthetic patterns via differences in growth rate using cells containing synthetic genetic edge detectors. **(A)** Different phases of the moon. Image of sample starting microcolonies on the left. Images of different phases of the moon. Growth rates of green fluorescing cells relative to sector edge detection cells are 1.1, 1.0, and 0.6 from left to right. (**B)** Different pacmans. Images of sample seeding (left) and grown pacmans with different sized mouths with BF image of colony inset. Growth rates of green fluorescing cells and non-fluorescing cells relative to sector edge detector cells are 1.1, 1.0, and 0.9 from left to right. **(C)** Yinyang-like patterns. Image of sample seeding (left) and grown yinyang-like colonies. Scale bars are 500 µm unless specified otherwise.

## DISCUSSION

### Performance limits of the border functions

We abstracted and developed border functions that guide the engineering of the curvature, texture, and reliability of cell-lineage borders. The border functions govern how border features arise as an emergent consequence of cell physical properties. Specifically, the *curvature* border function defines border endpoint angles achieved as a function of cell growth rates. The *texture* border function defines root mean square of borders from their average trajectories as a function of cell drag coefficient. The endpoint *reliability* border function defines the variation of achieving a specific border endpoint angle as a function of cell shape. These border functions, as now defined, can be used to more predictably engineer border curvature within a range of border endpoint angles of 30° to 120°, border texture with cells protruding from 10 to 60 µm in the opposite cell lineage, and border reliability with standard deviations of endpoint angles ranging from 15° to 23°.

### Abstracted functions highlight scientific opportunities to understand patterning

The border functions highlight opportunities for improving our understanding for how cells collectively realize precise patterning. The texture and reliability border functions suggest that spherical cells do not make the smoothest or most reliable patterns. Instead, short rod cells, cells with a small length-to-diameter aspect ratio produce the smoothest and most reliable patterns. In colloidal systems, short rod particles have been observed to achieve higher packing densities than perfect spheres (Donev et al., 2004; Williams and Philipse, 2003). Furthermore, systems with short-rod particles densely packed approach a glass transition while spherical and rod shaped cells do not (Bolhuis and Frenkel, 1997); tight packing of short rod shaped cells may cause a population of such cells to more readily “freeze” in place, preventing cell buckling and creating smoother and more reliable borders.

### Border functions serve as a language to enable use of a holistic approach to control border features

Practically, one barrier to utilizing holistic patterning in synthetic morphogenesis has been the complexity of pattern emergence as a consequence of underlying cell physical properties. Global patterns arise across space and time as a result of many cell level interactions that effect the whole. Even a relatively small biological system, such as a colony of *E. coli* one millimeter in radius, is made up of millions of cells. The force calculation that determines the position of each cell increases exponentially with the number of cells as every cell interacts with the whole colony, and the resulting microscopic pattern features are often non-intuitive.

### Abstraction hierarchy as a unifying framework for synthetic morphogenesis

To help manage the complexity of traversing between explicit synthetic genetic code and growth and formation of patterns, we offer an initial abstraction hierarchy that supports engineering of synthetic morphogenesis in a form compatible with an existing abstraction hierarchy (Figure 7) (Endy, 2005). Here, we focus specifically on allowing engineers to relate specific and tunable physical properties of cells that can be used to program borders, such as growth rate and cell shape, to the resulting growth of desired patterns. Stated differently, the border functions abstraction layer developed here give guidance for what cell operator values will grow a desired pattern; engineers wishing to realize a pattern may traverse this hierarchy via simple exchanges. The abstraction hierarchy also presents a common framework to facilitate a community effort for improving how to better engineer synthetic morphogenesis. For example, we hope the abstraction functions will be extended to not only include more cell-specific characteristics (e.g., adhesiveness and elasticity), but also cell signaling operators, cell motility, and differentiation.

**Figure 7:**
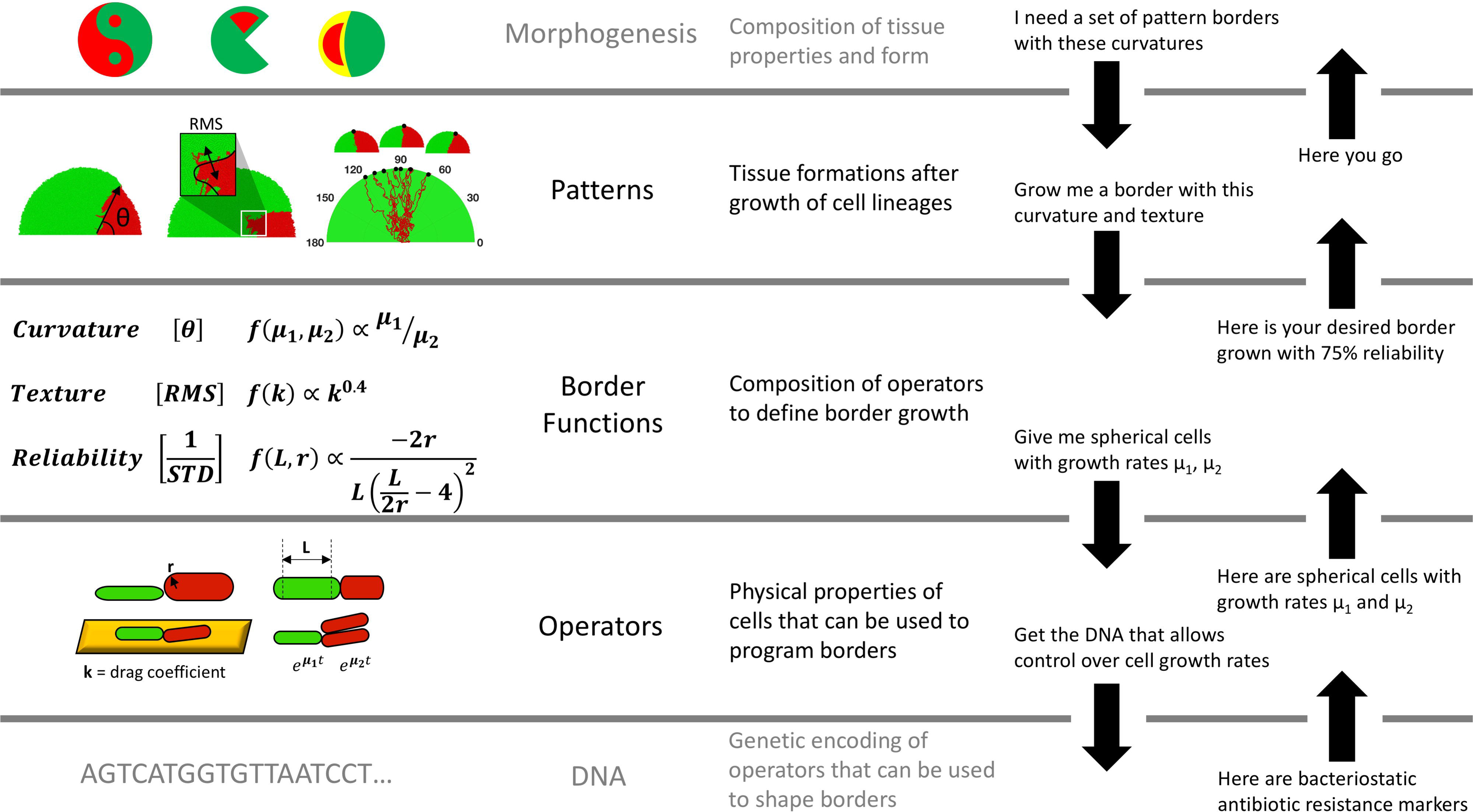
A proposed abstraction hierarchy supporting the holistic engineering of synthetic morphogenesis. The purpose of this abstraction hierarchy is to manage the complexity of growing synthetic biological patterns starting with DNA very similar to the abstraction hierarchy for building genetic systems. Efforts to engineer synthetic morphogenesis are divided into five layers. The first layer is the DNA that dictates a cell’s physical properties. The second layer is the cellular operators, or the physical properties of the cell such as cell growth rate and shape that effect the overall pattern. The third layer is the border functions layer that predicts border curvature, texture, and endpoint reliability based upon cell operators. The fourth layer are resulting patterns from the growth of cell lineages. The fifth layer are patterns resulting from the composition of patterns from the fourth layer.

### Towards engineering the growth of more complex biological patterns

Natural patterning systems realize complex morphogenesis via a variety of mechanisms. Past work understanding natural morphogenesis and engineering synthetic morphogenesis largely focused on genetic systems implementing reaction-diffusion mechanisms as the primary means of temporal and spatial patterning (Basu et al., 2005; Cooper et al., 2015; Danino et al., 2010; Gregor et al., 2007; Raspopovic et al., 2014; Skopelitis et al., 2017; Sohka et al., 2009; Tabor et al., 2009). However, there is an opportunity to take advantage of a holistic approach to patterning. Recent work harnessed individual cell characteristics to realize synthetic patterns and studies have started to focus on the mechanical basis of development (Cachat et al., 2016; Mori et al., 2009; Sampathkumar et al., 2014; Uyttewaal et al., 2012). A holistic approach that combines growth with synthetic genetic systems to forward engineer the growth, formation, and healing seems both necessary and sufficient to fully realize the power and understanding implied in mastery of synthetic morphogenesis.

## ACKNOWLEDGMENTS

We thank Eric Wei, Curt Frank, and Akshay Maheshwari for useful discussions, Rohinton Kamakaka for his guidance in micromanipulation, Fernan Federici for his help with plasmid segregation, Tim Rudge and Anton Kan for their help with CellModeller, and Jonathan Calles for his help editing. Funding was provided by the National Science Foundation Graduate Research Fellowship under Grant No. DGE-114747 and additional support was provided by Stanford University.

## AUTHOR CONTRIBUTIONS

AC and DE designed the experiments. AC carried out the experiments. AC and DE interpreted the data, and wrote and approved the manuscript.

