## Supplemental Information for "Abstracted functions for engineering the autonomous growth and formation of patterns"

**Figure S1: Estimation of curves in yinyang-like pattern**

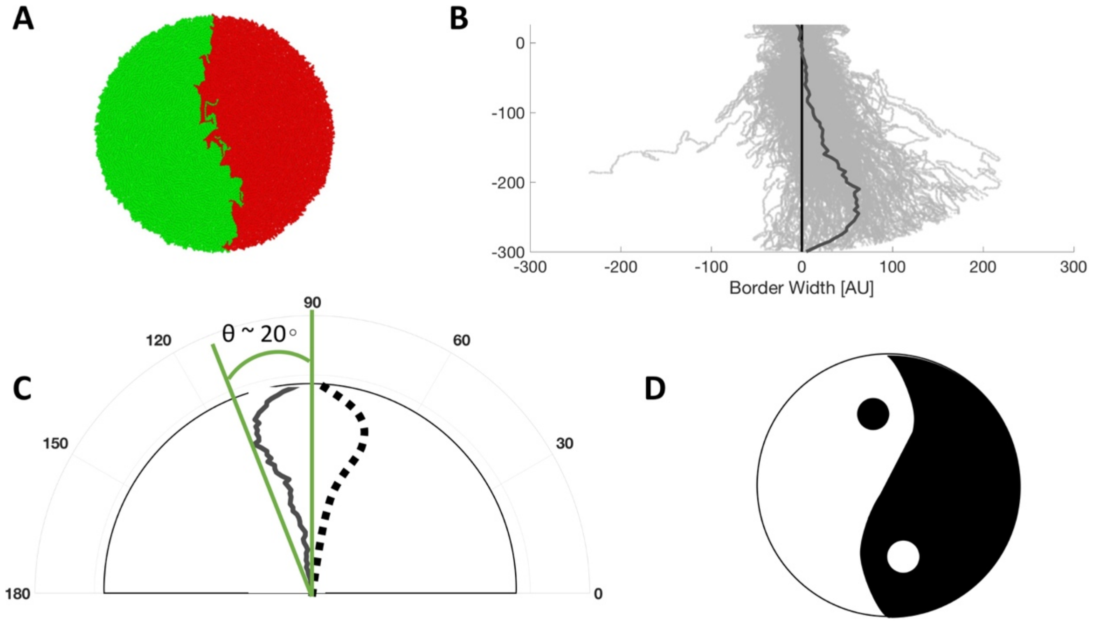

(A) A sample simulation of four cells growing at different growth rates creating the curves of a yinyang-like pattern.
(B) Borders of 100 simulations (gray traces) were averaged (dark gray line).
(C) The average border of a yinyang-like pattern does not create a bend as sharp as a yinyang.
(D) The average border applied to predict a yinyang-like pattern created by the envisioned development program.

**Figure S2: Sector edge detector system design and characterization**

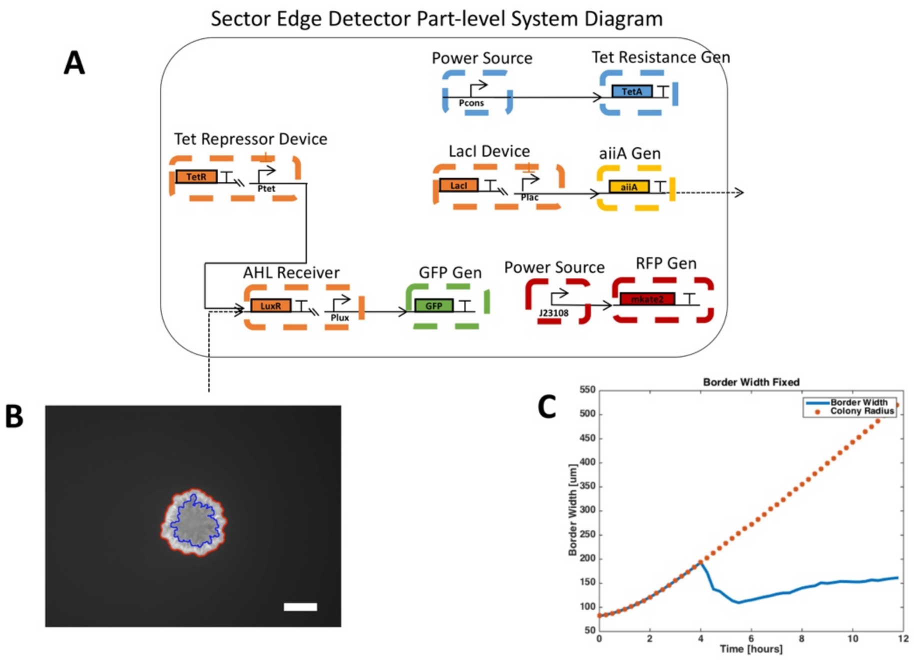

(A) Part-level system diagram of sector edge detector. Sector edge detector constitutively expresses mkate2 and aiiA gene which degrades AHL. AHL induces expression of GFP.
(B) When a colony grows on agar with a homogenous concentration of AHL, aiiA accumulates and degrades AHL faster. Eventually, the diffusion of AHL equilibrates with degradation rate and only the colony edge receives enough AHL to trigger GFP expression. We track colony radius (red) and border width (blue).
(C) Border width is fixed at 150 μm over time.

**Supplementary Figure 3: Failure modes of growing a yinyang-like pattern**

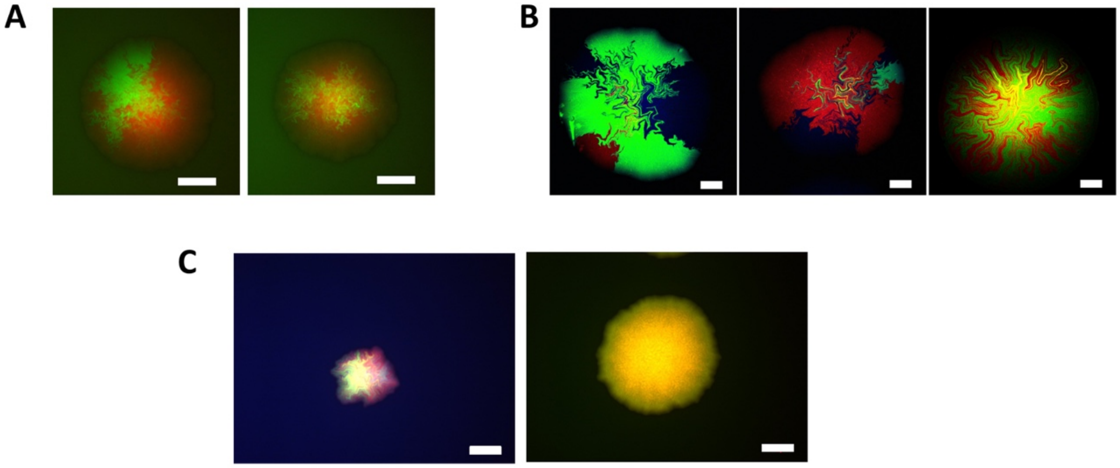

(A) Images of colonies grown for 16 hours from cells with two segregation plasmids. Multiple sectors signify multiple differentiation events during the growth of the colony.
(B) Images of colonies grown for 16 hours from cell with four segregation plasmids. Multiple sectors or less than four sectors indicate late or early segregation of four plasmids and that some colonies did not have all plasmids.
(C) Image of colony grown from cells with four segregation plasmids (left) indicates the segregation of all four plasmids by the presence of all four fluorescent signals in separate sectors of the colony. Image of colony grown with sector edge detection plasmids (right) indicates the sector edge detection plasmids do not segregate by the absence of unique fluorescence in different colony sectors. Scale bars 500 µm.

**Supplementary Figure 4: Border roughness increases during exponential growth of colony but does not increase during nutrient-depleted growth.**

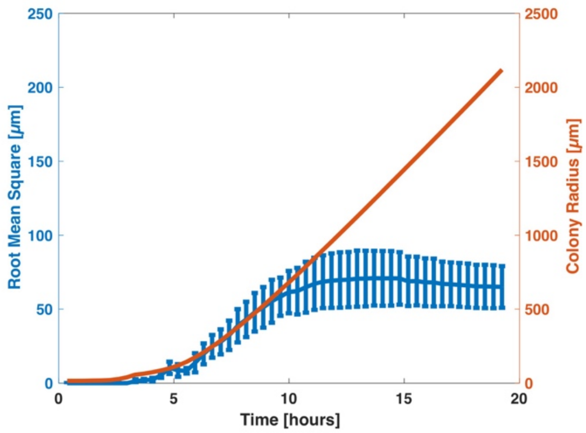

Simulated border roughness over time (blue) with a standard deviation above and below each point. Colony radius is plotted in orange. Border roughness is quantified as the root mean square between the border and a rolling average of the border. N = 100.

**Supplementary Figure 5: Endpoint angle is a measure of border curvature**

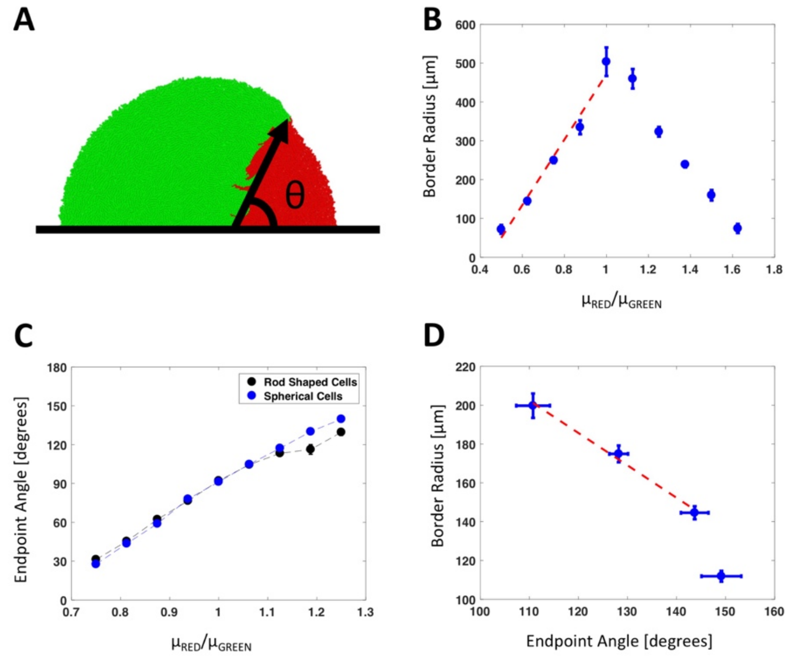

(A) Border endpoint angle is defined as the angle between the wall cells are growing against and the line from the beginning of the border to the end of the border.
(B) Mean border curvature radius depends linearly on the ratio of growth rates. Border curvature radius is obtained by taking the radius of a circle fitted to the border. Standard error of endpoint angle is plotted at each point. $R^2 = 0.9760$ N = 80.
(C) Mean border endpoint angle is plotted for different growth rates for spherical cells and rod shaped cells. Standard error of endpoint angle is plotted at each point. N = 100.
(D) Linear relation between border endpoint angle and border curvature radius in the range of endpoint angles examined in this work. $R^2 = 0.9915$ N = 100.

**Supplementary Figure 6: Increasing mechanical constraints leads to more reliable and smoother borders**

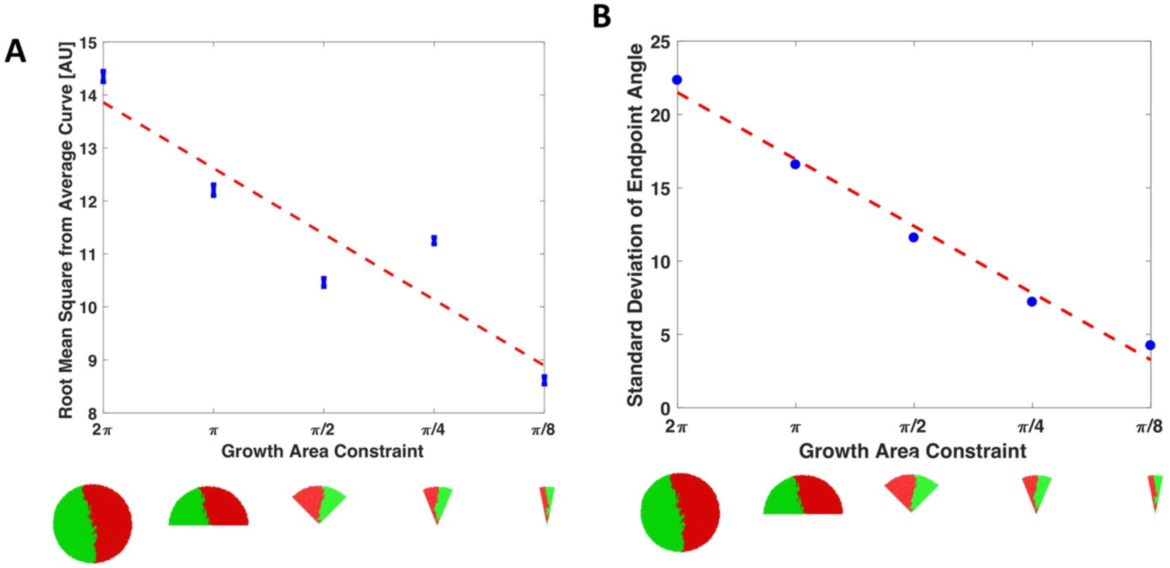

The growth area is gradually constrained from a total growth area angle of 2π all the way to an angle of π/8. Sample simulation results are shown below the x-axis.

(A) Border roughness is plotted for different growth area constraints with standard error plotted above and below each point. $R^2 = 0.8575$ N = 1000.

(B) Standard deviation of border endpoint angle is plotted for different growth area constraints with standard error plotted above and below each point. $R^2 = 0.9868$ N = 1000.

**Supplementary Figure 7: Variability in bucking and division length causes unreliable border growth.**

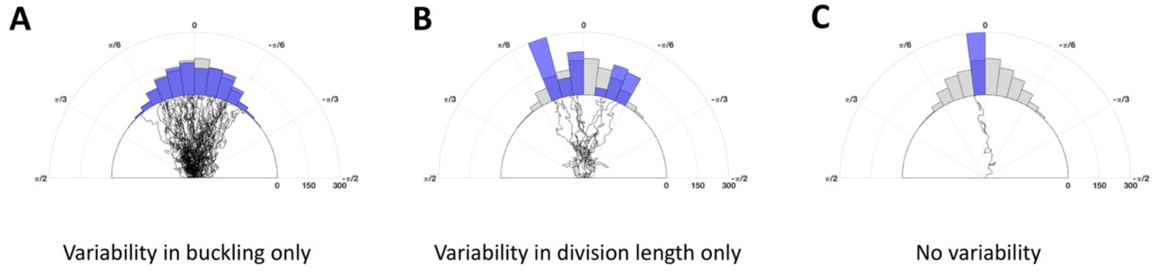

(A) Polar histogram of border endpoints from 1000 simulations with variability in cell buckling only in blue. Polar histogram of border endpoints from 100 simulations with variability in both cell division length and cell buckling in gray.
(B) Polar histogram of border endpoints from 1000 simulations with variability in cell division length only in blue.
(C) Polar histogram of border endpoints form 1000 simulations without variability in cell division length and cell buckling in blue.

**Supplementary Figure 8: Texture and reliability measurements are precise enough.**

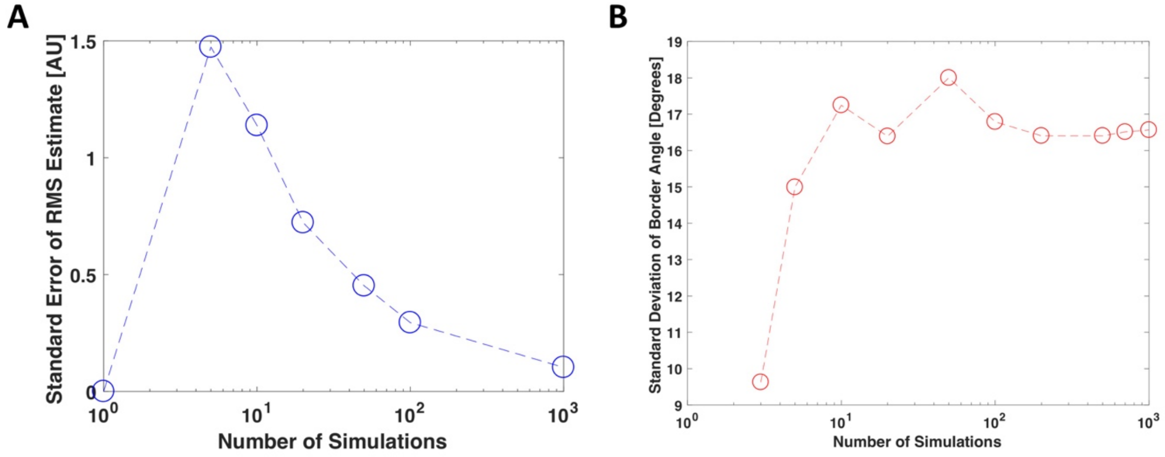

(A) Estimating root mean square of border from its rolling average from a sample size of N = 100 resulted in a value ~0.25 AU off of the population mean.
(B) Estimating standard deviation of border angle from a sample size of N = 100 results in an estimate less than 1/2 degree off of the population standard deviation.

**Supplementary Figure 9: Sensitivity analysis of critical border function operators**

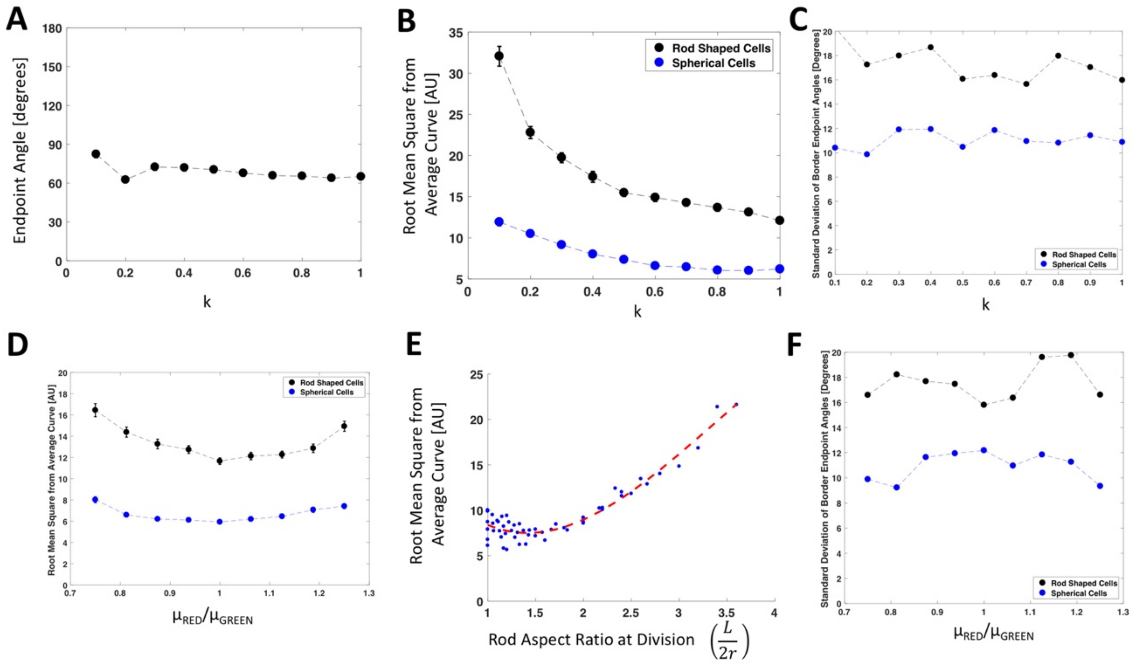

(A) Plot of mean endpoint angle for different drag coefficients per cell length. Simulations were run with a growth rate ratio of $\mu_{RED}/\mu_{GREEN} = 0.875$. Standard error of endpoint angle is plotted at each point.
(B) Average border root mean square from smooth border is plotted for different drag coefficients for both rod and spheres. Standard error is plotted at each data point.
(C) Standard deviation of endpoint angles resulting from growth on different drag coefficients per cell length are plotted for rods and spheres.
(D) Plot of mean border texture at different growth rate ratios for spherical and rod shaped cells. Standard error is plotted at each data point.
(E) Average border root mean square from smooth border is plotted for different rod aspect ratios at division.
(F) The standard deviation of endpoint angles is plotted at different growth rate ratios for rods and spheres.
Sample sizes for all data points is N = 100.

**Supplementary Figure 10: Micromanipulation seeding characterization**

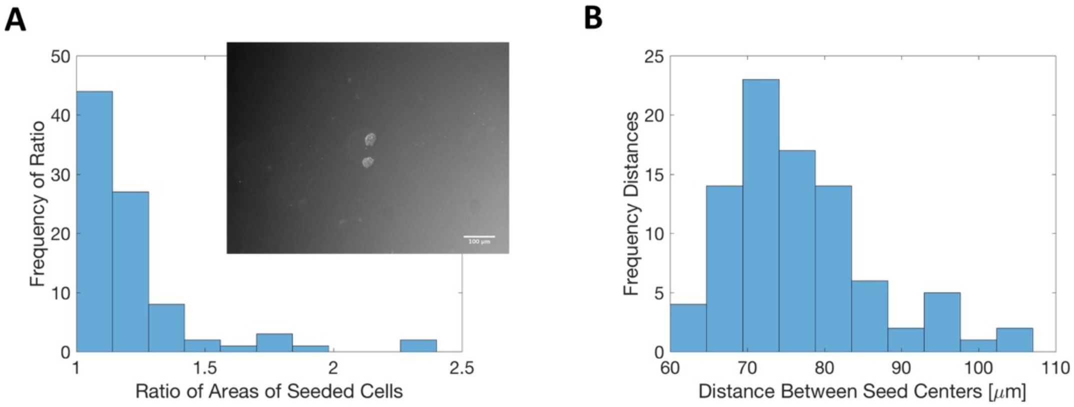

(A) Cell types were micromanipulated as microcolonies next to each other. Inset is a sample micromanipulation event. Histogram of area ratios of 100 micromanipulation events. If the ratio exceeded 1:1.2, the event was not used in border function experiments.
(B) Histogram of distances between microcolony centers of 100 micromanipulation events.

**Supplementary Figure 11: Maximum permissible ratio of microcolony areas**

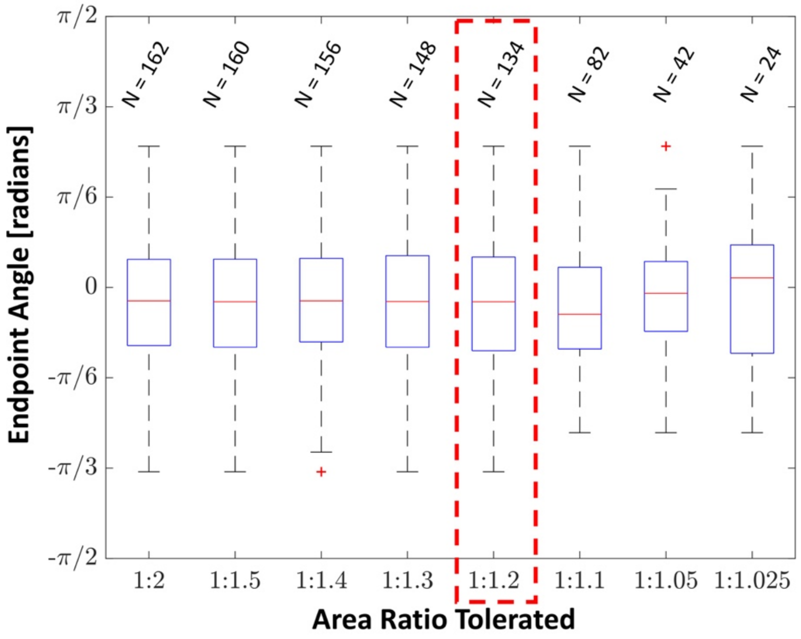

Minimization of area ratio tolerated while maximizing N to keep the average endpoint angle consistent. Box plots of endpoint angles from colonies grown of rod shaped cells. Distribution of endpoint angles at different ratios of microcolony areas were characterized. At a ratio of 1:1.2, the N is large enough to keep the endpoint angle consistent.

**Supplementary Figure 12: Modulating cell growth rates by titrating bacteriostatic antibiotics**

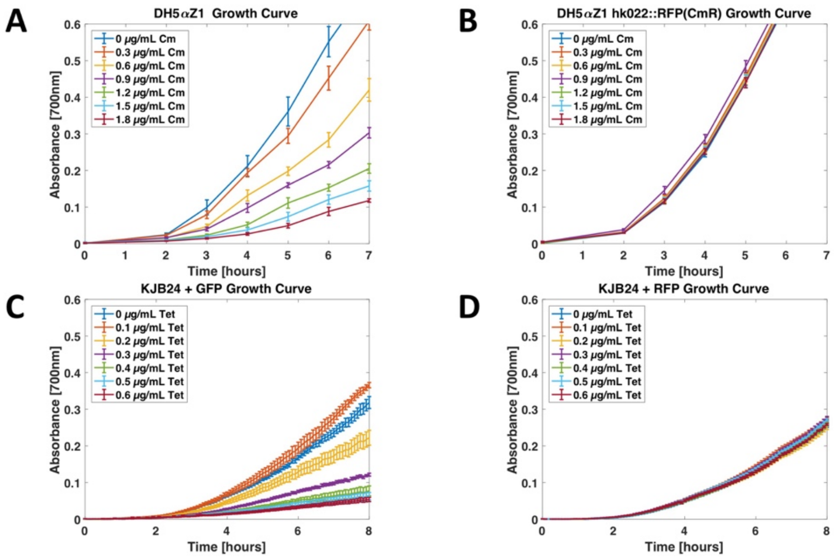

(A) Growth curves of rod shaped cells titrating chloramphenicol
(B) Growth curves of rod shaped cells with chloramphenicol resistance titrating chloramphenicol
(C) Growth curves of spherical cells titrating tetracycline
(D) Growth curves of spherical cells with tetracycline resistance titrating tetracycline

**Supplementary Figure 13: Morphology of cells is unaffected by antibiotic dosages used**

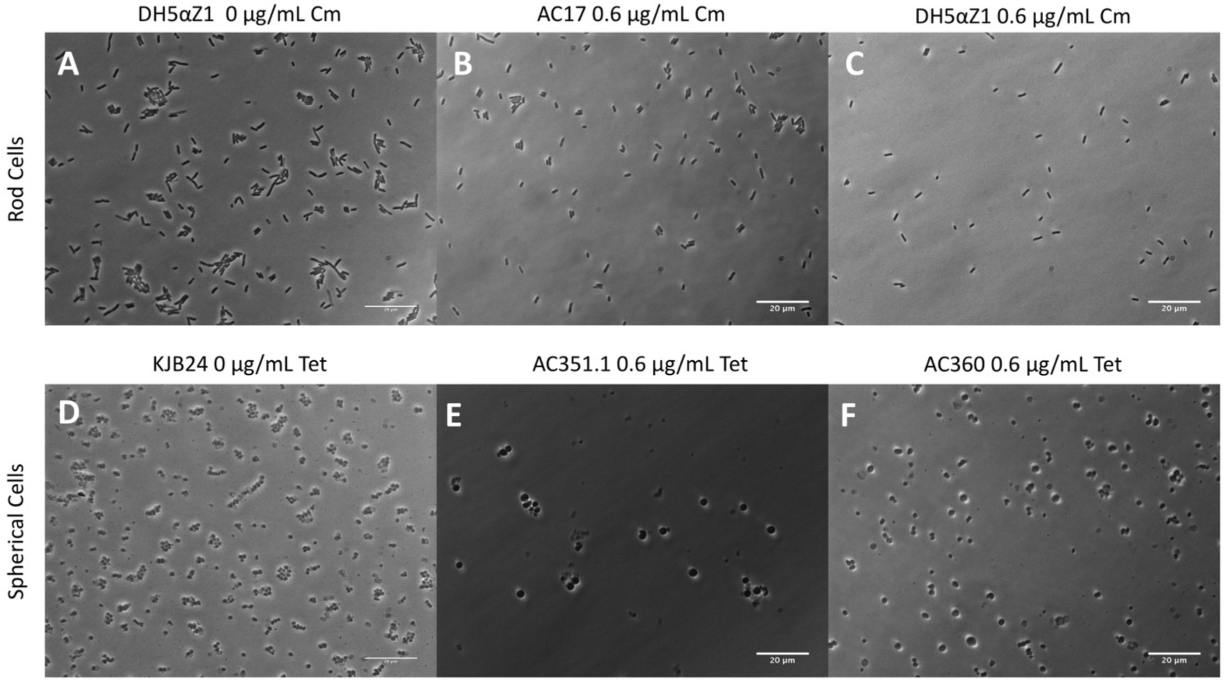

(A) 60X image of DH5αZ1 (rod shaped) cells without antibiotics.
(B) 60X image of DH5αZ1 cells resistant to chloramphenicol induced with the maximum dosage of chloramphenicol used in experiments.
(C) 60X image of DH5αZ1 cells not resistant to chloramphenicol induced with the maximum dosage of chloramphenicol used in experiments.
(D) 60X image of KJB24 cells (spherical) cells without antibiotics.
(E) 60X image of spherical cells not resistant to tetracycline in presence of tetracycline.
(F) 60X image of KJB24 cells resistant to tetracycline induced with the maximum dosage of tetracycline used in experiments.

**Supplementary Figure 14: Quantification of morphology of cells is unaffected by antibiotic dosages used**

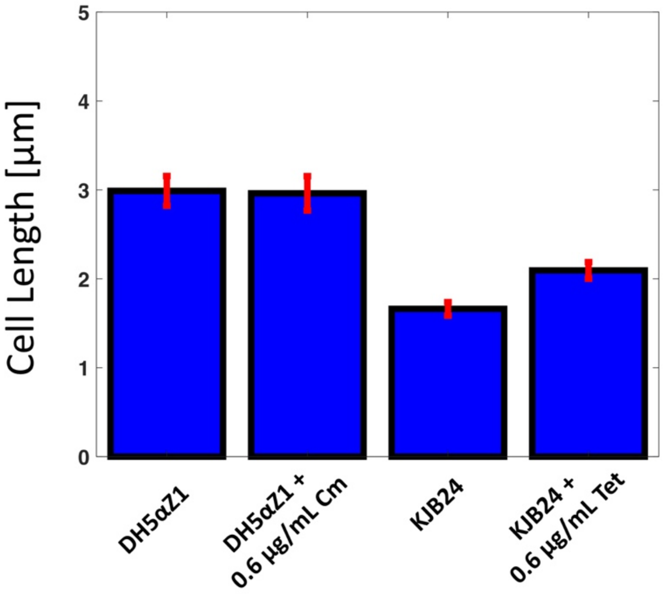

Mean DH5αZ1 cell lengths with and without maximum dosage of chloramphenicol used.. KJB24 cell diameter mean with and without maximum dosage of tetracycline used. Tetracycline increases the size of cells slightly. Standard error above and below the mean approach the limit of measurement accuracy of 0.1 µm

**Supplementary Figure 15: Differences between simulations and experiments**

| | **Simulations** | **Experiments** |
|---|---|---|
| Doubling Time | 34 minutes | ~66 minutes |
| **Time of Exponential Growth** | **0-9 hours** | **0-9 hours** |
| **Time of Nutrient Depleted Growth** | **9+ hours** | **9+ hours** |
| Number of Cells Seeded | 1 cell | ~100 cells |
| Area of Grown Colony | ~0.01 $mm^2$ | ~1 $mm^2$ |
| Initial Seeding Distance | 5 µm | ~75 µm |

We scale simulation parameters to match growth behavior observed in experiments in terms of time in exponential growth and time in nutrient depleted growth. Simulation results show that border roughness, reliability, and radius of curvature are largely determined during the exponential growth phase of colonies. At the same time, we scale down the number of cells to make the computation feasible.

**Supplementary Figure 16: Matching growth behavior of simulations and experiments**

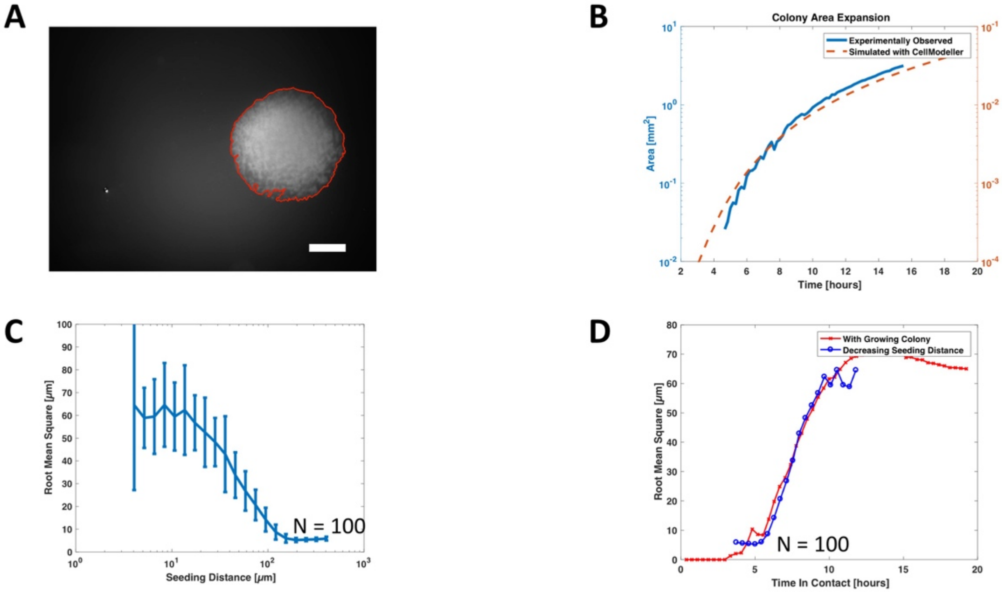

(A) Sample image of colony area tracking over time. Area tracked is enclosed in red line.
(B) Simulated and empirical growth behavior of colonies. Area of colony is roughly 100 times larger than simulated colony
(C) Border texture depends on cell seeding distance. Cells seeded further apart grow smoother borders.
(D) As cells are seeded further apart, cell lineages have less time in contact and less time for borders to become rough. Seeding distance is converted to time cell lineages are in contact with each other. Time cell lineages grow in contact with each other is a critical determinant of border texture.

**Supplementary Figure 17: Growth curves of strains used to grow different phases of the moon and pacmans**

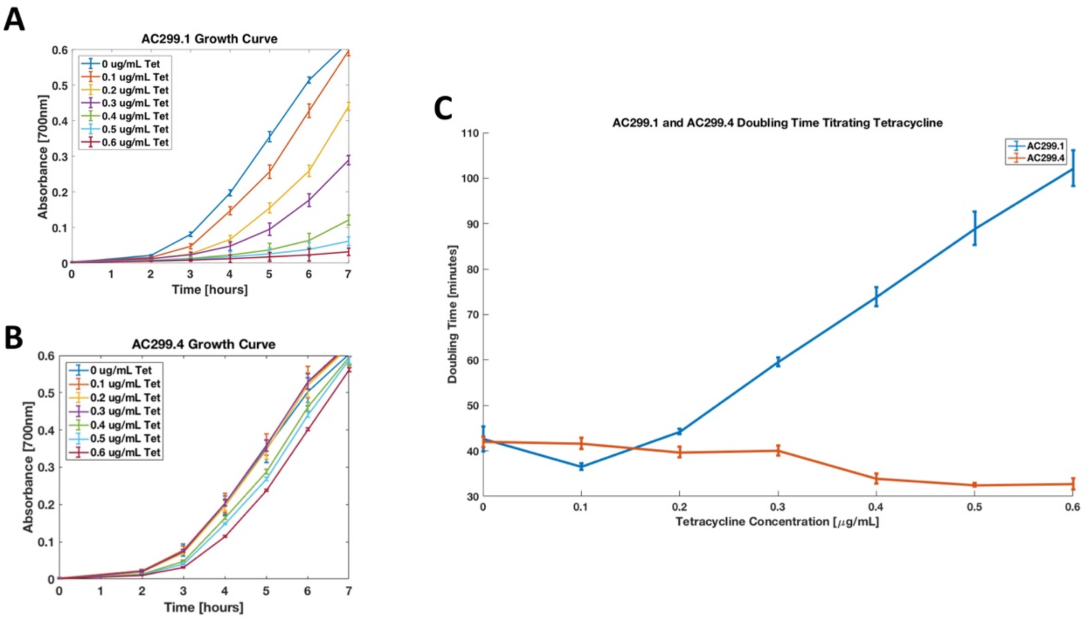

(A) Growth curves of DH5αZ1 cells expressing green fluorescence (AC299.1) titrating tetracycline
(B) Growth curves of DH5αZ1 harboring sector edge detection system (AC299.4) titrating tetracycline
(C) Doubling times of AC299.1 and AC299.4 plotted against tetracycline concentration. N = 3.

**Supplementary Figure 18: Growth curves of strains used to grow yinyang-like patterns**

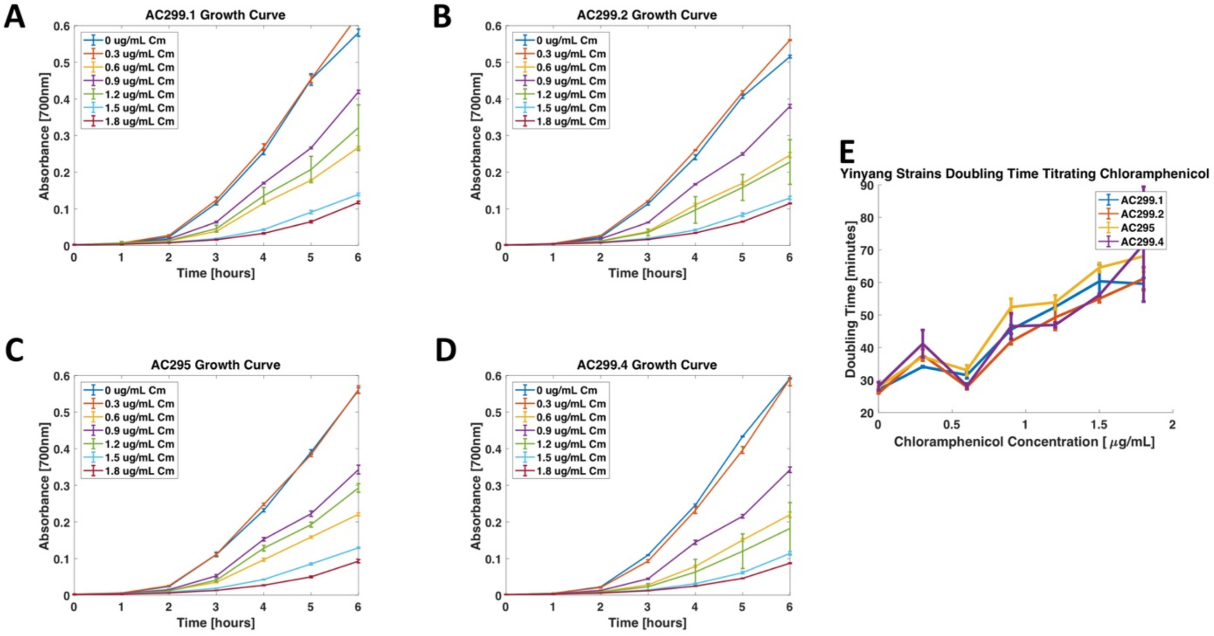

(A) Growth curves of DH5αZ1 cells expressing green fluorescence (AC299.1) titrating chloramphenicol

(B) Growth curves of DH5αZ1 cells expressing red fluorescence (AC299.2) titrating chloramphenicol

(C) Growth curves of DH5αZ1 harboring inverted sector edge detection system (AC295) titrating chloramphenicol

(D) Growth curves of DH5αZ1 harboring sector edge detection system (AC299.4) titrating chloramphenicol

(E) Doubling times of AC299.1, AC299.2, AC295, and AC299.4 titrating chloramphenicol

**Supplementary Figure 19: More images of grown yinyang-like colonies**

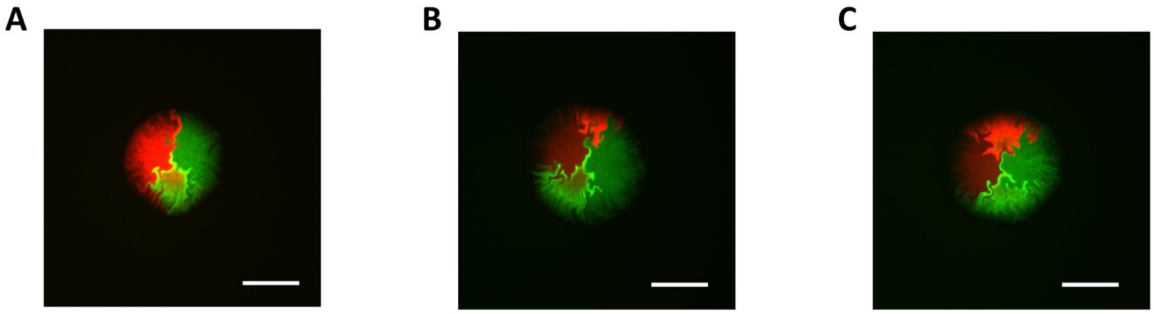

(A) Half a yinyang-like colony. A sample colony with the expected border curve of sector edge detector cells. The red sector edge detector cells got outcompeted.
(B and C) Other sample yinyang-like colonies.

**Supplementary Figure 20: Spherical cells grow a yinyang-like pattern more smoothly and reliably**

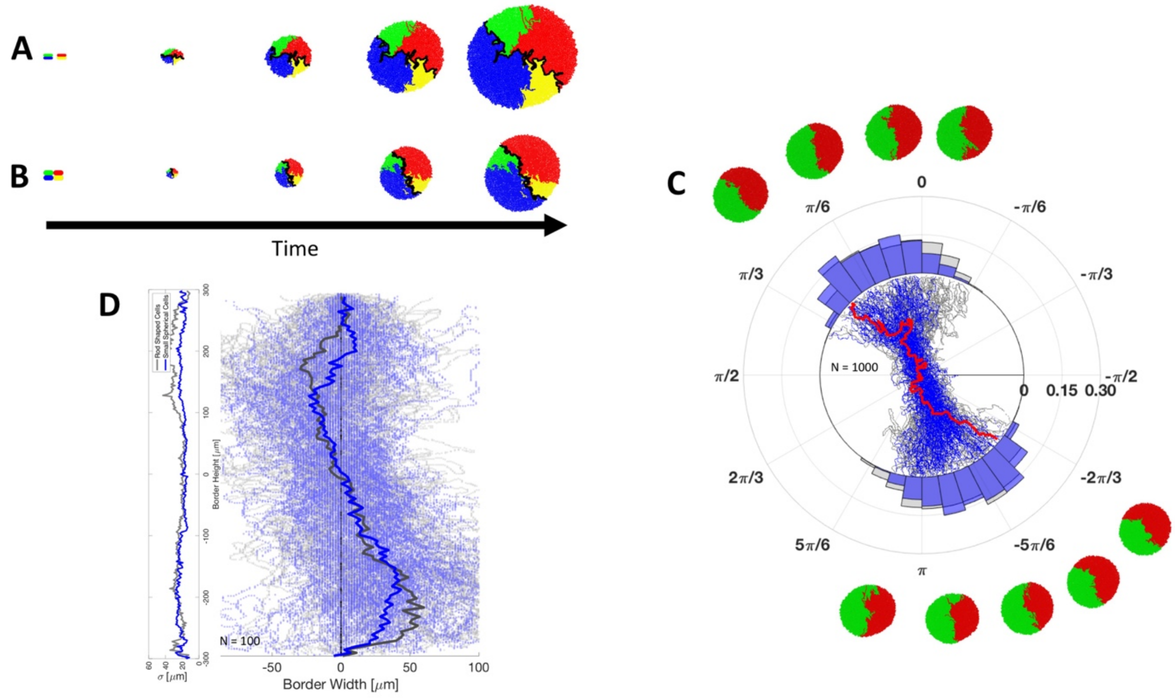

(A) Time-lapse images of yinyang-like pattern growth simulation with rod shaped cells. Yinyang-like pattern border is highlighted in black. The first image is enlarged to show how cells are initially seeded.

(B) Time-lapse images of yinyang-like pattern growth simulation with spherical cells.

(C) Spherical cells grow a yinyang-like pattern more reliably than rod shaped cells. Polar histogram of border endpoint angles of yinyang-like patterns grown from rod shaped cells (gray) and spherical cells (blue). Sample patterns grown from spherical cells are shown above and sample patterns grown from rod shaped cells are shown below.

(D) Spherical cells grow a yinyang-like pattern with smoother borders than rod shaped cells. Borders of patterns are fixed at the endpoints to observe border texture. The average border grown by spherical cells is traced in blue while the average border grown by rod shaped is traced in gray. At each border height, the standard deviation of border traces is calculated and plotted at the left.

**Supplementary Figure 21: Digital image processing algorithm used to analyze pattern borders**

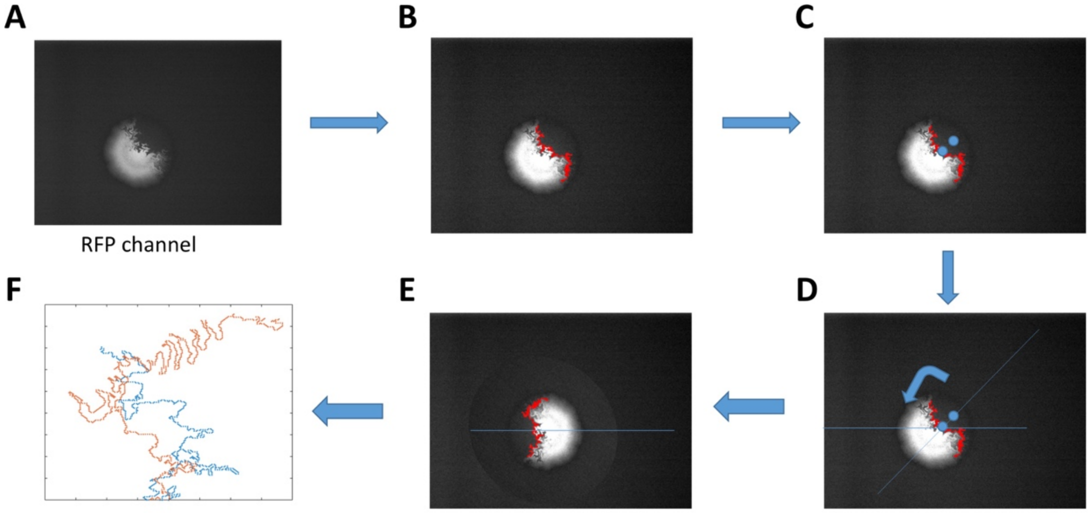

(A) The colony is first imaged in the BF, RFP, and GFP channel.
(B) Border is found by an "AND NOT" operation of the edge detected in the RFP channel and edge detected in the BF channel.
(C) The center of the border is found by minimizing the distance of points on the border to the colony center. The center of the smallest sector is found.
(D) The image is rotated to align the line connecting the middle of the border and the center of the smaller sector and the line at 180°.
(E) The border is then split into two borders.
(F) The border center is aligned at (0,0) and the bottom border is reflected about the 180° line.

#### Appendix 1: Script used for CellModeller simulations

```
import random
from CellModeller.Regulation.ModuleRegulator import ModuleRegulator
from CellModeller.Biophysics.BacterialModels.CLBacterium import CLBacterium
from CellModeller.GUI import Renderers
import numpy
import math

cell_cols = {0:[0,1.0,0], 1:[1.0,0,0]}
cell_lens = {0:2.5, 1:2.5}
cell_growr = {0:0.8, 1:0.8}
max_cells = 10000;
def setup(sim, GR):
    # Set biophysics, signalling, and regulation models
    biophys = CLBacterium(sim, max_cells=max_cells, jitter_z=False, muA=GR[3])
    biophys.addPlane((0,0,0), (0,1,0), 1.0)

    global cell_growr
    cell_growr = {0:0.8,1:GR[0]}
    global cell_lens
    cell_lens = {0:GR[2], 1:GR[2]}

    # use this file for reg too
    regul = ModuleRegulator(sim, sim.moduleName)
    # Only biophys and regulation
    sim.init(biophys, regul, None, None)
    #sim.outputDirName = '~/'

    # Specify the initial cell and its location in the simulation
    sim.addCell(cellType=0, length=GR[2], pos=(-2.5,0.5,0), dir=(1,0,0), rad=GR[1])
    sim.addCell(cellType=1, length=GR[2], pos=(2.5,0.5,0), dir=(-1,0,0), rad=GR[1])
    sim.pickleSteps = 10
    sim.outputSteps = 10

def init(cell):
    cell.targetVol = cell_lens[cell.cellType] + random.uniform(0.0,0.5)
    cell.growthRate = cell_growr[cell.cellType]
    cell.color = cell_cols[cell.cellType]

def update(cells):
    #Iterate through each cell and flag cells that reach target size for division
    for (id, cell) in cells.iteritems():
        if cell.volume > cell.targetVol:
            cell.divideFlag = True

def divide(parent, d1, d2):
    # Specify target cell size that triggers cell division
    d1.targetVol = cell_lens[parent.cellType]
    d2.targetVol = cell_lens[parent.cellType]
```

**Appendix 2: Matlab Scripts used for CellModeller on AWS**

2A. Parent script used to create simulation commands, get instance IP addresses, copy files to instances, run simulations, and download simulations results:

```
function inst_ips = GA_AWSinterface4(Curr_Pop,gen_indx,num_inst,time)

%A. Create runScripts
runScripts = AWS_ScriptCreate2(Curr_Pop,num_inst);
a = 'done with A'
fclose all;

% B. Get ip addresses
inst_ips = AWS_getIps;
a = 'done with B'

% C. Copy files to instances
for inst_indx = 1:length(inst_ips)
   if gen_indx == 1

AWS_ScriptCopy(inst_ips(inst_indx),["CLBacterium.py";"char_sim3.py";"Simulator2.py";"batch_four.py";runScri
pts(inst_indx)])
   else
      AWS_ScriptCopy2(inst_ips(inst_indx),runScripts(inst_indx))
   end
end
a = 'done with C'

% D. Run simulations
for inst_indx = 1:length(inst_ips)
   AWS_Clean(inst_ips(inst_indx));
   AWS_runSimulations(inst_ips(inst_indx),runScripts(inst_indx));
end
a = 'done with D'

% E. Download images and clean memory
unix(['mkdir /Users/atrichoksi/cellmodeller/data/EC2/ToyEvol/gen' int2str(gen_indx)]);
for inst_indx = 1:length(inst_ips)
   AWS_Download(inst_ips(inst_indx),gen_indx);
   AWS_Clean(inst_ips(inst_indx));
end
a = 'done with E'
```

2B. Creating run scripts

```
function runScripts = AWS_ScriptCreate2(array,num_inst)

%Atri Choksi
%March 27, 2017
%Creating scripts to copy over to instances and run
Pop_size = size(array,2);
```

```
for indx = 1:num_inst
    runScripts(indx) = strcat("runScript",int2str(indx));
    indx2 = (Pop_size*(indx-1)/num_inst + 1):(Pop_size*indx/num_inst);
    fid = fopen(['/Users/atrichoksi/Documents/Programming/MATLAB/EvolutionaryAlgorithms/AWS_Scripts/'
char(runScripts(indx))],'wt');

    template =
fileread('/Users/atrichoksi/Documents/Programming/MATLAB/EvolutionaryAlgorithms/AWS_Scripts/runScript_te
mp_char.txt');

    fprintf(fid,template,array(:,indx2));

    unix(['chmod u+x /Users/atrichoksi/Documents/Programming/MATLAB/EvolutionaryAlgorithms/AWS_Scripts/'
char(runScripts(indx))]);
end
```

#### 2C. Get IP address

```
function inst_ips = AWS_getIps

[~,inst_ips] = unix('aws ec2 describe-instances | grep -i PublicIpAddress  | awk "{ print $2}" | cut -d"," -f1| sed -e
"s/"//g"');
inst_ips = strrep(inst_ips,'.','-');
inst_ips = strsplit(string(inst_ips),'\n');
inst_ips(end) = [];
end
```

#### 2D. Copy scripts to instances

```
function AWS_ScriptCopy2(IP,filenames)

%Atri Choksi
%March 28, 2017
%To copy files specified in filename to the instance with ip address IP

cd /Users/atrichoksi/AmazonWebServices/AWS_Scripts
KNAME = 'AWSEC2keypair.pem';
for indx = 1:length(filenames)
    unix(['scp -o StrictHostKeyChecking=no -i ' KNAME ...
        ' /Users/atrichoksi/Documents/Programming/MATLAB/EvolutionaryAlgorithms/AWS_Scripts/'
char(filenames(indx))...
        ' ec2-user@ec2-' char(IP) '.us-west-2.compute.amazonaws.com:/home/ec2-user/CellModeller/Scripts'])
end
```

#### 2E. Running simulations on instances

```
function AWS_runSimulations(IP,scriptname)

%Atri Choksi
%March 28, 2017
%Runs cellmodeller on instance

cd /Users/atrichoksi/AmazonWebServices/AWS_Scripts
```

```
KNAME = 'AWSEC2keypair.pem';
unix(['ssh -i ' KNAME ' ec2-user@ec2-' char(IP) '.us-west-2.compute.amazonaws.com -f /home/ec2-user/CellModeller/Scripts/' char(scriptname)])
end
```

#### 2F. Downloading simulations results

```
function AWS_Download(IP,generation)

%Atri Choksi
%March 28, 2017

cd /Users/atrichoksi/AmazonWebServices/AWS_Scripts
KNAME = 'AWSEC2keypair.pem';

unix(['mkdir /Users/atrichoksi/cellmodeller/data/EC2/ToyEvol/gen' int2str(generation) '/' char(IP)]);
unix(['scp -i ' KNAME ' ec2-user@ec2-' char(IP) '.us-west-2.compute.amazonaws.com:/home/ec2-user/cellmodeller/data/*.jpg /Users/atrichoksi/cellmodeller/data/EC2/ToyEvol/gen' int2str(generation) '/' char(IP)])
%unix(['ssh -i ' KNAME ' ec2-user@ec2-' char(IP) '.us-west-2.compute.amazonaws.com -f "rm -rf /home/ec2-user/cellmodeller/data/*"'])
```

#### Appendix 3: Cells and Plasmids Used

| Name | Host Strain | Plasmid | Description |
|---|---|---|---|
| DH5αZ1 | DH5αZ1 | - | Cloning strain of *Escherichia Coli* |
| AC17 | DH5αZ1 hk022::RFP($Cm^R$) | - | Rod shaped cells expressing RFP. |
| AC19 | DH5αZ1 hk025::RFP($Kan^R$) | - | Rod shaped cells expressing RFP. Growth rate modulatable with chloramphenicol. |
| AC299.1 | DH5αZ1 | BAC GFP, $Carb^R$, $Kan^R$ | GFP expressing cells used for figure 6. |
| AC299.2 | DH5αZ1 | BAC mkate2, $Carb^R$, $Zeo^R$ | RFP expressing cells used for figure 6. |
| AC295 | DH5αZ1 | BAC GFP, $Carb^R$, $Kan^R$, aiiA, LuxR, $P_{lux}$ mkate2 | Cells that express GFP everywhere and mkate2 at edges of sector |
| AC299.4 | DH5αZ1 | BAC mkate2, $Carb^R$, $Tet^R$, aiiA, LuxR, $P_{lux}$ GFP | Cells that express mkate2 everywhere and GFP at the edges of sector |
| KJB24 | W3110 rodA(Am) ddlB::Tn5 | | Spherical cells obtained from the Haseloff lab |
| AC351.1 | KJB24 | (SEG10) GFP, $Cm^R$, $Carb^R$ | Spherical cells expressing GFP. Growth rate modulatable with tetracycline |
| AC360 | KJB24 | (SEG11) mCherry, $Cm^R$, $Tet^R$ | Spherical cells expressing RFP. Growth rate not affected by tetracycline. |

### Appendix 4: Plasmid Maps

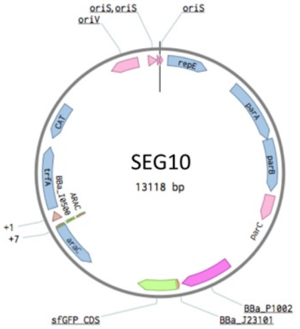

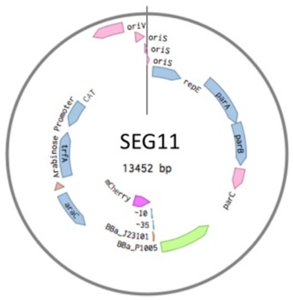

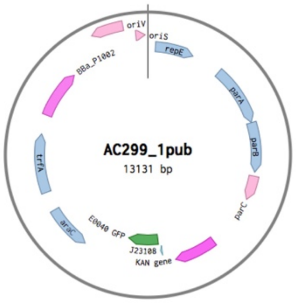

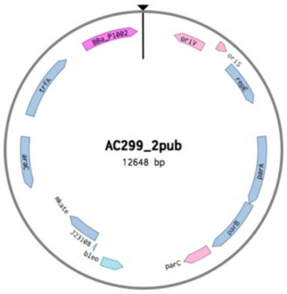

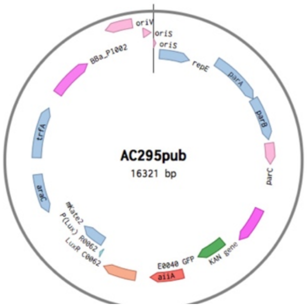

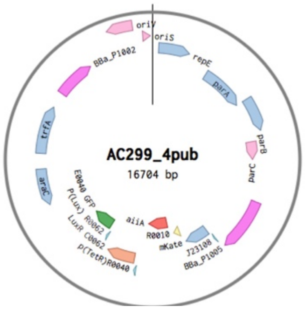

### Appendix 5: Primers used

| Primer Name | Sequence 5' – 3' |
|---|---|
| ACp145 | CCACTCATCGCAGTATTACGTAGCAATCAACTCACTGGC |
| ACp146 | GAACTTTCAGCGGTACTAAATACATTCAAATATCTATCCGCTCATGAG |
| ACp272 | gccctgtcaggggcggggttttttttttctagtattattatttgtatagttcatcc |
| ACp283 | tttacagctagGTTTAAACctgacagctagctcagtcctaggtataatgctagcaTctactagagtcacacaggaaagtac |
| ACp274 | CGGGCTGAGGTATCGATACCTTAGCCtaaaaaaaaccccgccctgtcaggggcg |
| ACp285 | acaggaaagtactagatgaaattatatatggaaggtactgtcaacaatc |
| ACp284 | gccctgtcaggggcggggttttttttttctagtattattatcaacgatgtcctaatttcgacgg |
| ACp225 | GGAGGTTTTCTAatgacagtaaagaaactttatttcatcccagc |
| ACp226 | ctatcagagttattaatattccgggaacactctacaactc |
| ACp229 | GGCTAAGGTATCGATACCTCAGCCCGGCACATAGAAGTAAcaatacgcaaaccgcctctcc |
| ACp230 | TGGAGTTTTGGCGCGCCAGAGTGACACAGTCCTTaaacgcagaaaggcccacc |
| ACp169 | tatagtcgaataaaTctactagagtcacacaggaaagtactag |
| ACp170 | cagcgagtcagtgagtataaacgcagaaaggcccac |
| ACp113 | TAAGGTATCGATACCTCAGCC |
| ACp116 | ggagttgtggtaatctatgtatcc |
| ACp150 | TTACTTCTATGTGCCGGGC |
| Acp185 | gtttAAGGACTGTGTCACTCTGG |
| ACp341 | AGCTAACTTACATTAATTGCGTTGC |
| ACp342 | gccatgatgtatacattgtg |
| ACp346 | ttgcgagaatgtcaaacgc |
