## Supplementary Materials for "Abstracted functions for engineering the autonomous growth and formation of patterns"

### 1. Simulations

CellModeller is a GPU based biofilm growth simulator. CellModeller version 4.2.1 was used for all simulations (*1*). MATLAB v2017A was used to coordinate the run of CellModeller in Amazon Web Service (AWS) instances (The MathWorks Inc., USA, Amazon Web Services, USA).

#### 1.1 Simulation Conditions

A fixed wall was placed at (0,0) extending in the x direction. Initial cells were placed against each other parallel to the wall right above the origin. A target length for cell division was set before and cells divided when they reached that target volume plus a random number from a uniform distribution between 0 and 0.5 $\mu$m. Simulations were run until the number of cells exceeded 10,000. Yinyang-like pattern simulations were run under similar conditions except the fixed wall was removed. CellModeller's inbuilt CLBacterium.py script was used to simulate the physics. Appendix 1 contains the python script used to run the simulations.

The same initial conditions produced different simulation results due to randomness in cell buckling and cell division length. Randomness in cell buckling was implemented as described in the original Cellmodeller publication (*1*). Randomness in cell division was implemented by sampling a random uniform distribution between 0 and 0.5 $\mu$m and adding it to the target cell length for division of each cell.

#### 1.2 Simulating on Amazon Web Services

p2 instances on AWS were rented. CellModeller was installed in a disk image with a preinstalled NVIDIA CUDA Toolkit 7.5 on Amazon Linux. MATLAB scripts were used to copy CellModeller python scripts to AWS along with a shell script that ran CellModeller in AWS (Appendix 2). Resulting colony images were downloaded.

### 2. Molecular Biology

2.1 Cell used

Cells used in this study are listed in appendix 3. DH5αZ1cells were used for all rod shaped cell experiments and pattern growth experiments. KJB24 cells obtained from the Hasseloff lab were used for all experiments involving spherical cells (*2*).

2.2 Plasmid Backbones

Bacterial artificial chromosome (BAC) backbones were obtained from Fernan Federici (*3*, *4*). BACs with carbenecillin resistance (SEG10), tetracycline resistance (SEG11), spectinomycin resistance (SEG12), and kanamycin resistance (SEG13) were received. SEG 10 constitutively expressed GFP under J23101 promoter and had chloramphenicol and carbenecillin resistance. The second BAC, SEG11, constitutively expressed mCherry and had chloramphenicol and tetracycline resistance.

2.3 Cloning materials and methods

All cloning PCR reactions were performed with 2X Phusion Mastermix (ThermoFisher Scientific, USA) using a 30sec/kb extension time. All DNA assembly reactions were performed with Infusion Reaction Mastermix (Clontech,USA). Primers and gblocks were purchased from IDT. All cells and plasmids used in this work are listed in appendix 3. Plasmid maps are shown in appendix 4. Primers are given in appendix 5.

2.4 Yinyang Plasmid Construction

AC299.1 was constructed by first replacing the chloramphenicol resistance gene with a ampicillin resistance gene in SEG13 creating SEG14. The ampicillin resistance cassette was cloned from SEG10 using PCR with primers ACp145 and ACp146. Next EGFP with a J23108 promoter was PCR amplified twice from plasmid #1009 from Bonnet et al. 2012 using primers ACp272 and ACp283 followed by ACp274 and Acp283 and inserted in after the kanamycin resistance gene (*5*).

AC299.2 was constructed by replacing kanamycin resistance gene in SEG14 with a zeomycin resistance gene gblock. Next, mKate2 under a J23108 promoter was PCR amplified with two consecutive PCR reactions from plasmid #1009 using primers ACp285 and ACp284 followed by ACp283 and ACp274.

AC295 was constructed by cloning Ptet LuxR, Plac aiiA, and Plux mkate2 into AC299.1. Ptet LuxR and Plux mkate2 were purchased as gblocks, and Plac aiiA was PCR amplified off of pTD103aiiA from the Hasty lab using primers ACp225 and ACp226 followed by ACp229 and ACp230 on the PCR product (*6*).

AC299.4 was constructed by replacing chloramphenicol resistance cassette in SEG11 with carbenicillin resistance in the same way as done to construct AC299.1. Next, J23108 mkate2 was sequentially PCR amplified from plasmid #1009 with primers ACp285 and ACp284 followed by ACp283 and ACp274. Plux GFP was PCR amplified from plasmid #1009 with primers ACp169 and ACp170. Ptet LuxR and Plac aiiA were obtained in the same way as done to construct AC295.

Plasmids were sequence with primers ACp113, ACp116, ACp150, ACp185, ACp341, ACp342, and ACp346 to confirm plasmid inserts.

**3. Growth Curves**

Growth curves by measuring optical density at 700nm to avoid interference from RFP (*7*). The SpectraMaxI3 plate reader was used to take reading every hour. Here is the step-by-step procedure used to obtain the growth curves:

a. Streak glycerol stocks of cells on LB agar plate
b. Grow cultures overnight in 5 mL LB at 37C shaking at 270 RPM
c. Setting up plate and taking measurements
    i. Dilute cultures to O.D. 1
    ii. Add 198 $\mu$L of LB with 1% Arabinose + 1000 ng/mL aTc, 1M IPTG, and 1$\mu$M AHL + antibiotic to each well in a 96 well plate
    iii. Add 2 $\mu$L of dilute culture to each well
    iv. Take t = 0 reading of O.D. 700
    v. Shake plate at 37C
    vi. Transfer plate to platereader and take OD 700 every hour

**4. Experimental Implementation of Border Function Experiments**

For rod shape cell experiments, DH5αZ1 cells were seeded next to AC17 (DH5αZ1 hk025::RFP(CmR)) and grown for 10 hours and 40 minutes at 37C before imaging. Cells were seeded on LB agar plates with or without sublethal doses of chloramphenicol to modulate growth rate of DH5αZ1 cells.

KJB24 cells with SEG10 were seeded next to KJB24 cells with SEG11 for spherical cell experiments. Cells were seeded and grown in the same way as rod shaped cells were except tetracycline was used to modulate growth rates and cells were allowed to grow for 16 hours.

Cells were seeded with the following protocol:

a. Streak glycerol stocks of cells on LB agar plate
b. Grow cultures overnight in 5 mL LB at 37C shaking at 270 RPM
c. Make LB Agar plates + sublethal doses of Cm
d. Backdilute cells 1:100 next morning
e. Allow growth to 0.4 O.D
f. Drip streak cultures at 10x concentrated cells.
g. Make sure you let the plates dry enough at a neutral angle
h. Plate 10 uLs
i. Micromanipulate two cell types next to each other
j. Micromanipulate colonies with 10 mm spacing

**5. Pattern Growth Experiments**

To grow different phases of the moon, AC299.4 cells were micromanipulated next to AC299.1 cells. Tetracycline was used to modulate the growth rate of AC299.1 cells. 0 $\mu$g/mL tetracycline was used for to grow the smallest moon, 0.1 $\mu$g/mL tetracycline was used to grow the second moon, and 0.3 $\mu$g/mL tetracycline was used to grow the last moon. Cells were seeded on LB agar plates with 1% Arabinose, 1000 ng/mL aTc, 1M IPTG, and 1$\mu$M AHL.

To grow different pacmans, AC299.1, AC19 (DH5αZ1 hk025::RFP(KanR)), and AC299.4 were seeded next to each other. Tetracycline was used to modulate the growth rate of AC299.1 and AC17. The pacman with the widest mouth was seeded on 0.14 $\mu$g/mL tetracycline, the pacman with a medium sized mouth was grown on 0.16 $\mu$g/mL tetracycline, and the pacman with the closed mouth was grown on 0.2 $\mu$g/mL tetracycline. Cells were seeded on LB agar plates with 1% Arabinose, 1000 ng/mL aTc, 1M IPTG, and 1$\mu$M AHL.

To grow yinyang-like patterns, AC299.1, AC299.2, AC295, and AC299.4 cells were seeded next to eachother. 0.9 $\mu$g/mL Chloramphenicol was used to equilibrate growth rates of the four strains. Cells were seeded on LB agar plates with 1% Arabinose, 1000 ng/mL aTc, 1M IPTG, and 1$\mu$M AHL.

**6. Imaging**

A Nikon TE2000 inverted microscope was used for imaging. 2X objectives were used for images of colonies and 10X microscope was used for imaging seeded cells. GFP was excited with a LambdaXL xenon arc light through a ET470/40X filter and fluorescence through a ET525/50 filter was collected. mKate2 was excited through a ET572/35X filter. The resulting fluorescence was collected through a ET632/60 filter. The light was captured using a coolsnapHQ2 camera. The images were analyzed using ImageJ.

### 7. Digital Image Processing

MATLAB was used to analyze both simulated and experimental images.

#### 7.1 Border endpoint angle

First, cell lineage borders were detected by finding the common edge between the green and red channels. Then, the border was transformed into points in Cartesian coordinates with the start of the border normalized to the origin. The points were sorted by either the x or y coordinate. Border endpoint was calculated by transforming the last point in the border into polar coordinates and taking the angle coordinate.

#### 7.2 Root mean square of border from a smooth curve

Once the border was detected and normalized and transformed into Cartesian coordinates, a Savitzky-Golay filter was used on the border to generate a smooth version of the border. The root mean square between the smooth border and the original border was calculated.

#### 7.3 Additional Processing On Experimental Images

Images of colonies in experiment posed the additional challenge of colony rotation. The colony had to be split into two hemispheres for analysis. The additional image processing is described in supporting figure 20.

### 8. Similarities and differences from simulations to experiment

We showed that spherical cells grow smoother and more reliable borders compared to rod shaped cells using both simulation and experiment. Although simulation and experimental conditions were not identical, we scaled conditions in order to keep the time of exponential growth and nutrient-depleted growth the same to match pattern growth behavior as closely as possible (Figure S19-20). Reliability of border endpoints in simulation and experiment differed by 6° and 4° for rod and spherical cells respectively. We hypothesize that standard deviations measured in simulations were consistently smaller than those in experiment because a mechanical constraint was applied to colonies in simulations (Figure S5). Roughness of spherical cells between simulation and experiment are similar, but roughness of rod shaped cells is three times larger in experiment than in simulation. Applying a high-pass filter to the experimental borders to subtract average border trajectories results in a border roughness similar to those observed in simulations. Similar border curvatures were achieved in simulations and experiments except when red fluorescing spherical cells were growing at 135% of the rate of green fluorescing spherical cells. We hypothesize this difference is because cells induced with antibiotics may reach a nutrient-depleted growth behavior more quickly.
